# Anterior thalamic function is required for spatial coding in the subiculum and is necessary for spatial memory

**DOI:** 10.1101/2020.01.31.928762

**Authors:** Bethany E. Frost, Sean K. Martin, Matheus Cafalchio, Md Nurul Islam, John P. Aggleton, Shane M. O’Mara

## Abstract

Hippocampal function relies on the anterior thalamic nuclei, but the reasons remain poorly understood. While anterior thalamic lesions disrupt parahippocampal spatial signalling, their impact on the subiculum is unknown, despite the importance of this area for hippocampal networks. We recorded subicular cells in rats with either permanent (*N*-methyl-D-aspartic acid) or reversible (muscimol) anterior thalamic lesions. Bayesian and other statistical analyses underscored the notable absence of the diverse spatial signals normally found in the subiculum, including place cells, following permanent anterior thalamic lesions. Likewise, there was marked disruption of these diverse spatial signals during transient lesions. By contrast, permanent anterior thalamic lesions had no discernible impact on CA1 place fields. Anterior thalamic lesions reduced spatial alternation performance (permanently or reversibly) to chance, while leaving a non-spatial recognition memory task unaffected. These findings, which help explain why anterior thalamic damage is so deleterious for spatial memory, cast a new spotlight on the importance of subiculum function and reveal its dependence on anterior thalamic signalling.

**Graphical Abstract:** 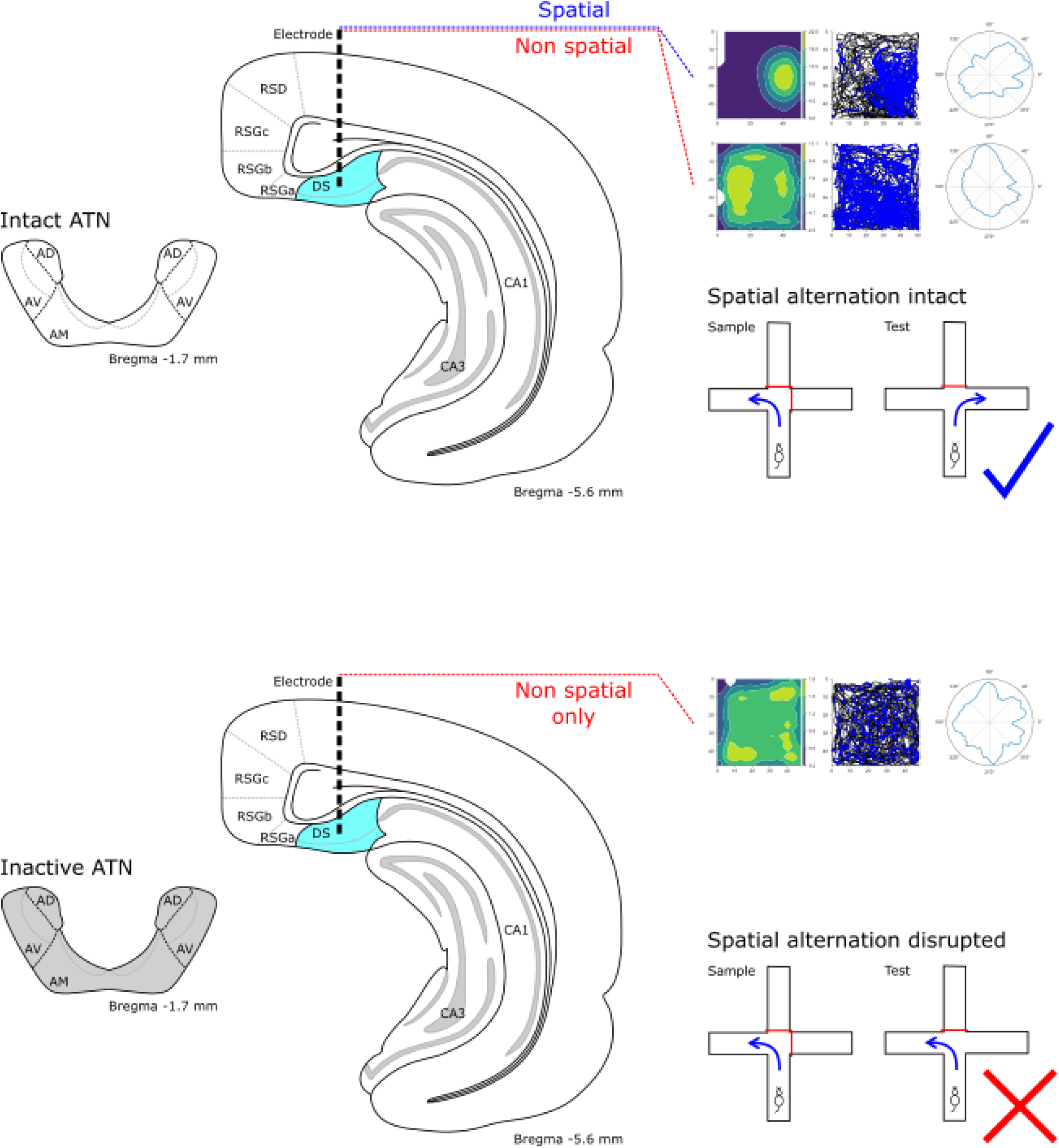

## Introduction

The hippocampal formation lies at the heart of the brain’s cognitive mapping capabilities^1^, while also comprising part of an extended system vital for human episodic memory^2–4^. The anterior thalamic nuclei (ATN), which consist of the anteromedial, anteroventral, and anterodorsal nuclei, are key structures within both functional systems^2,5^. In rodents, for instance, anterior thalamic lesions severely impair spatial learning, mirroring hippocampal damage^6–9^; moreover, disconnection studies point to the functional interdependence of the hippocampal formation and anterior thalamic nuclei^10–13^. Neuronal recordings within the ATN show the presence of place, head-direction, and border cells^14–16^, suggesting the ATN is also a spatial processing node.

Anterior thalamic lesions involving the anterodorsal nucleus eliminate parahippocampal head-direction signals^17,18^, while reducing the periodicity and frequency of parahippocampal grid cell activity^18^. Nevertheless, disruption of the head-direction network does not explain the severity or diversity of the spatial learning deficits following ATN lesions^19–22^. For example, head-direction system lesions (including the lateral mammillary nucleus, which provides head-direction signals for the ATN^23^), cause appreciably milder spatial deficits than ATN lesions^24–28^. The conclusion follows that the ATN contribute in important, substantive, and additional ways to spatial processing^21,29–30^.

Our attention has focussed on the subiculum, the principal hippocampal recipient of projections from the ATN^31,32^. The dorsal subiculum receives direct inputs from the anteroventral nucleus^31^, and is indirectly linked with ATN efferents via the the entorhinal cortices and retrosplenial cortex^33^. The direct thalamo-hippocampal inputs are, however, selective as CA1 does not receive ATN projections^31^. The dorsal subiculum contains cells with a variety of spatial properties, including place and grid cells, as well as boundary vector cells^34–37^. The relationship of these subiculum cell-types to the ATN remains unknown, though it might readily be assumed that the dense inputs from area CA1, with its numerous place cells^38,39^ are sufficient for effective subicular place cells. Likewise, inputs from entorhinal cortex^39^ could potentially ensure subicular grid cells. We, therefore, tested these assumptions by determining whether the spatial properties of subiculum cells depend on ATN activity^40,18^.

Here, we found lesions targeting the entire ATN (but sparing nucleus reuniens) abrogate subicular spatial signalling, and also reduce spatial alternation memory performance to chance. Bayesian and other statistical analyses underscored the notable absence of spatial firing in lesioned animals compared to controls. These analyses, thereby, emphasise the importance of ATN inputs for subicular spatial coding and their relevance for spatial alternation memory performance^10^. We then tested the anatomical specificity of this disconnection effect by recording hippocampal CA1 place cells after ATN lesions. The very dense CA1 projections to the subiculum^41–43^, which potentially supply the subiculum with spatial information^44^, provided the rationale for determining the sensitivity of CA1 to ATN damage^40^. Unexpectedly, we found that, although spatial alternation performance was again at chance after ATN lesions, place cell discharge in CA1 appeared largely unaffected.

## Results

Throughout, the terms hippocampal formation and hippocampal refer to the dentate gyrus, the CA fields, and the subiculum. The parahippocampal region includes the presubiculum, parasubiculum, postsubiculum and entorhinal cortices. Rats received permanent neurotoxic lesions of the anterior thalamic nuclei (ATNx) following injections of *N*-methyl-D-aspartic acid (NMDA), or transient ATN lesions following infusion of muscimol. Other rats served as controls.

### Combining anterior thalamic nuclei lesions (NMDA) with implantations in the dorsal subiculum

Of 23 animals implanted, one ATNx rat was subsequently excluded due to electrode malfunction, and one normal control was excluded because of post-surgical complications. Comparisons between the normal control and sham control groups consistently failed to reveal group differences and, for this reason, they were typically combined (group ‘CControl’).

### N-methyl-D-aspartic acid injections caused considerable cell loss within the anterior thalamic nuclei, but spared nucleus reuniens

Lesion effectiveness was quantified by comparing the total anti-NeuN reacted cell counts in the anteromedial, anterodorsal, and anteroventral thalamic nuclei in ATNx animals with the corresponding CControl values (Fig. 1A, B). The ATNx rats had markedly reduced cell counts (CControl 16209 ± 2507, ATNx 3497 ± 1528; t = −9.68, df = 8.26, p < 0.001, Welch’s Two Sample *t*-test; Fig. 1E), while the calbindin-reacted sections helped to confirm that nucleus reuniens remained intact (Fig. 1C, D), indicating that, despite a significant role in spatial working memory^45^, nucleus reuniens damage was not a contributing factor to differences in either behaviour or cell firing properties in ATNx animals. Maximum and minimum extent of the ATN lesions are shown in supplementary Fig. 1.

**Fig. 1:**
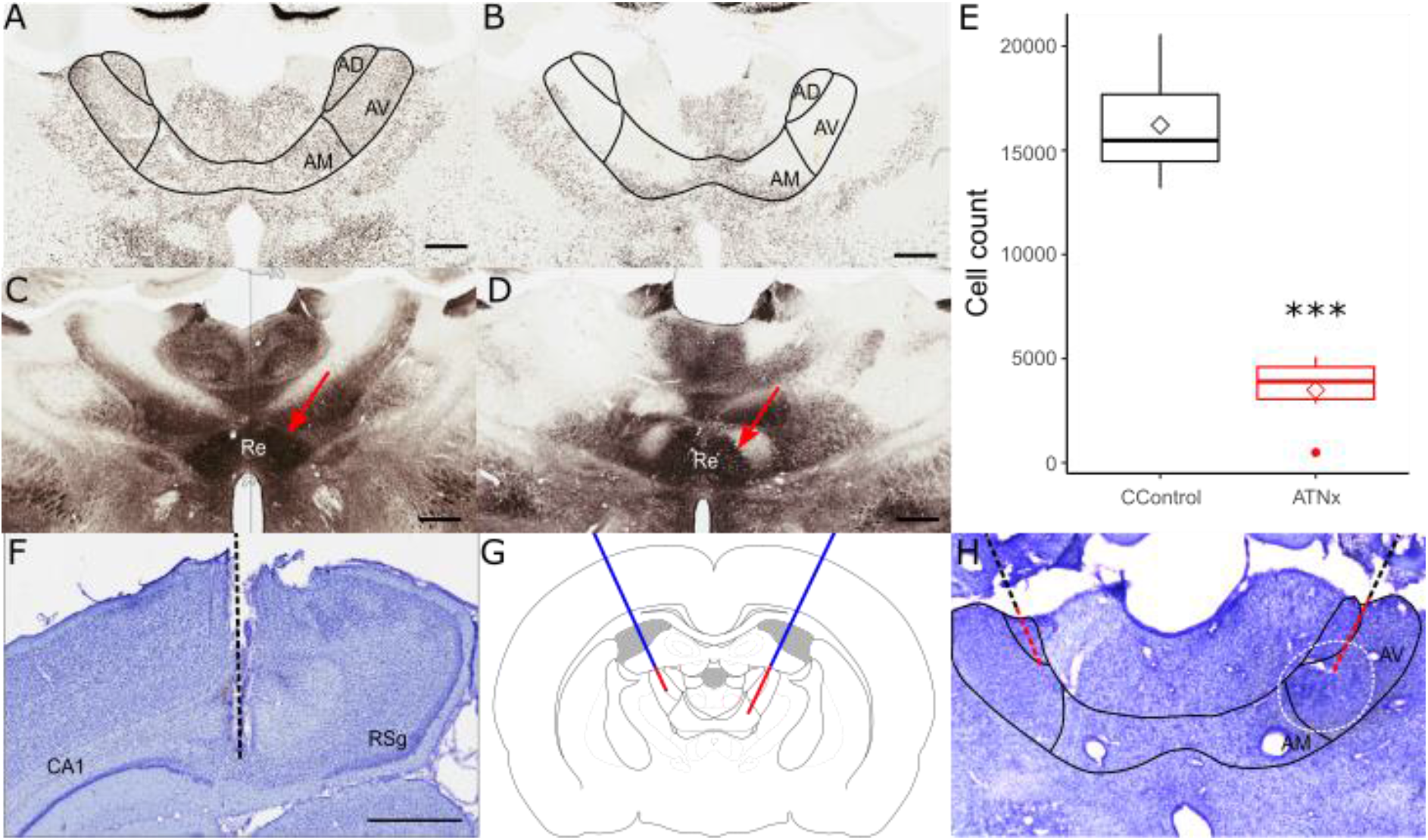
NeuN-reacted coronal sections showing the status of the anterior thalamic nuclei (ATN) in control (A) and lesion (B) animals. The nucleus reuniens (arrowed), as shown using calbindin-reacted sections, was intact in both control (C) and lesion (D) animals, indicating that reuniens damage was not responsible for deficits seen in ATN lesioned animals. (E) The ATN cell count was significantly reduced in lesioned animals (ATNx) compared to controls (CControl). Nissl-stained coronal sections helped to confirm electrode placement in dorsal subiculum (F), with the electrode path indicated. (G) Schematic representing cannula placement (blue) and the two targets of the infusion needle (red). (H) Cresyl violet stained section indicating cannula placement, with DAB-reacted flurogold infused to indicate spread of muscimol, with the black line indicating canula placement and red indicating the track of the infusion needle, and dashed white to indicate the spread of the muscimol. *** = p<0.001 (Welch’s Two Sample *t*-test). Scale bar = 800μm.

### ATN lesions (NMDA) reduce spatial alternation memory to chance levels of performance

Unless otherwise stated, Welch’s Two Sample *t*-tests were performed to compare CControl and lesion data.

Consistent with previous lesion studies, ATNx animals had considerably lower spatial alternation scores (52.23 ± 14.58%), compared to Normal (83.10 ± 10.96%) and Sham (81.25 ± 11.41%) controls (ANOVA p < 0.001, Tukey post-hoc p < 0.001; Fig. 2B), confirming their severely impaired spatial working memory. In contrast, ATNx animals showed no impairment in novel object discrimination when compared to the CControl animals (p > 0.05), indicating that recognition memory under these conditions remains intact (Fig. 2D-F). During free exploration in the square arenas with electrophysiological recordings, the ATNx rats travelled greater distances than the CControl animals both during habituation (CControl 62.14 ± 17.43 m, ATNx 85.45 ± 14.21 m; t = 3.55, df = 16.14, p = 0.003) and during subsequent recordings (CControl 120.66 ± 15.19 m, ATNx 142.41 ± 10.89 m; t = 2.98, df = 14.55, p = 0.096), suggesting slightly increased motor activity post-ATNx.

**Fig. 2:**
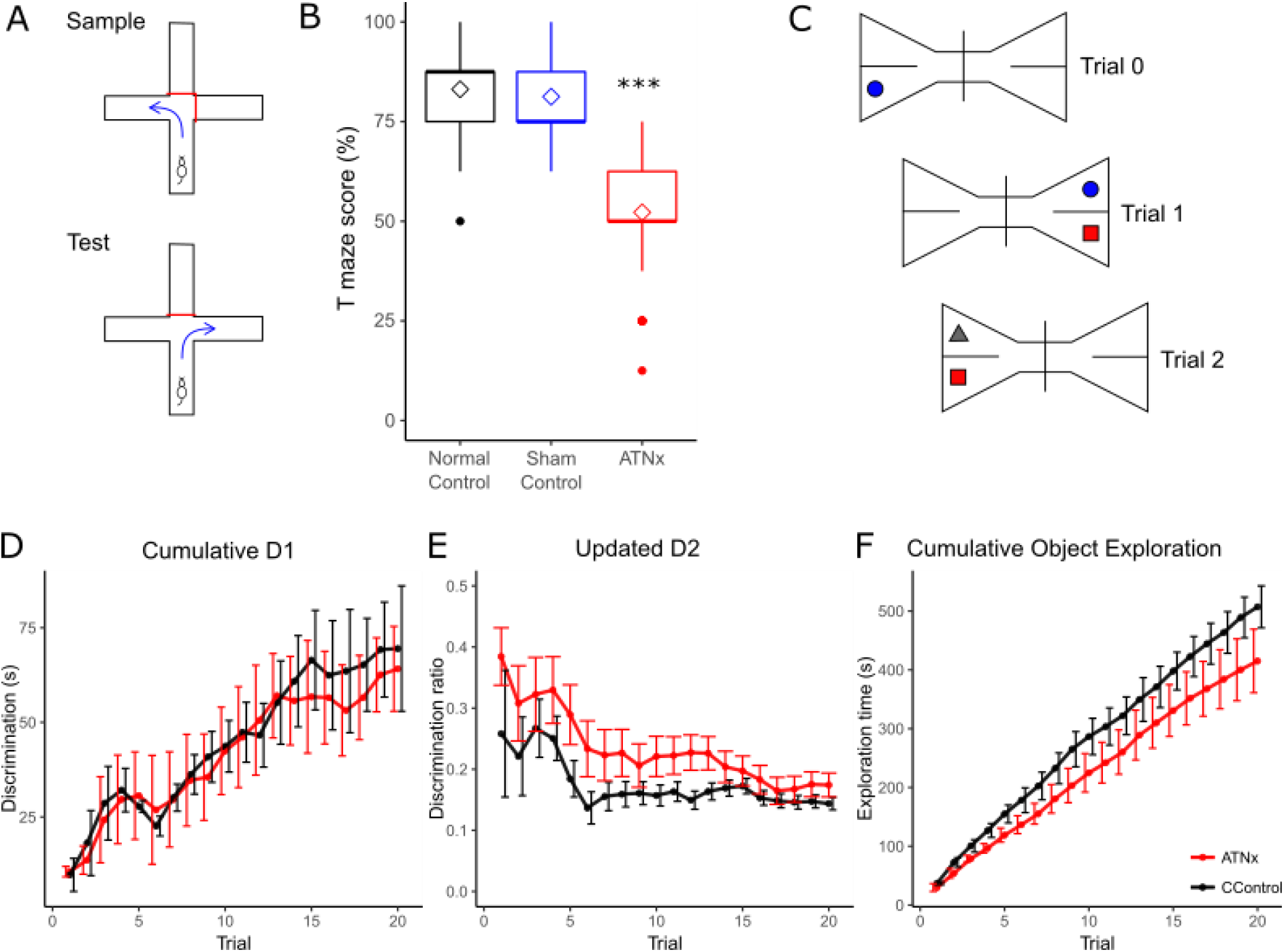
(A) Schematic diagram of spatial alternation task. ATN lesioned animals (ATNx) showed a significant deficit in spatial alternation compared to both control and sham animals (B). (C) Schematic of novel object recognition task. There was no difference between control (CControl) and lesion rats in cumulative D1 (D), D2 (E), or total exploration time (F). *** p<0.001 (ANOVA with Tukey post-hoc), error bars represent SEM.

### ATN lesions (NMDA) abrogate spatial firing in the subiculum only: Quantitative analyses

Spatial and non-spatial single units were recorded in the dorsal subiculum of all animals. Of 82 single units recorded in the CControl rats, 47 (57%) were considered spatial units (Fig. 3A). These units consisted of place (n = 11; 23%), head-direction (n = 20; 43%), border (n = 5; 11%) and grid (n = 11; 23%) cells. A further 35 (43%) did not show obvious spatial properties, e.g. no clear place field or preferred head direction. Strikingly, no spatial units were recorded in the ATNx animals, although non-spatial units (21), i.e., those showing no preferential firing in specific place or head orientation, were present (Fig. 3B). Electrode tracks are reconstructed in Fig. 1F-H.

**Fig. 3:**
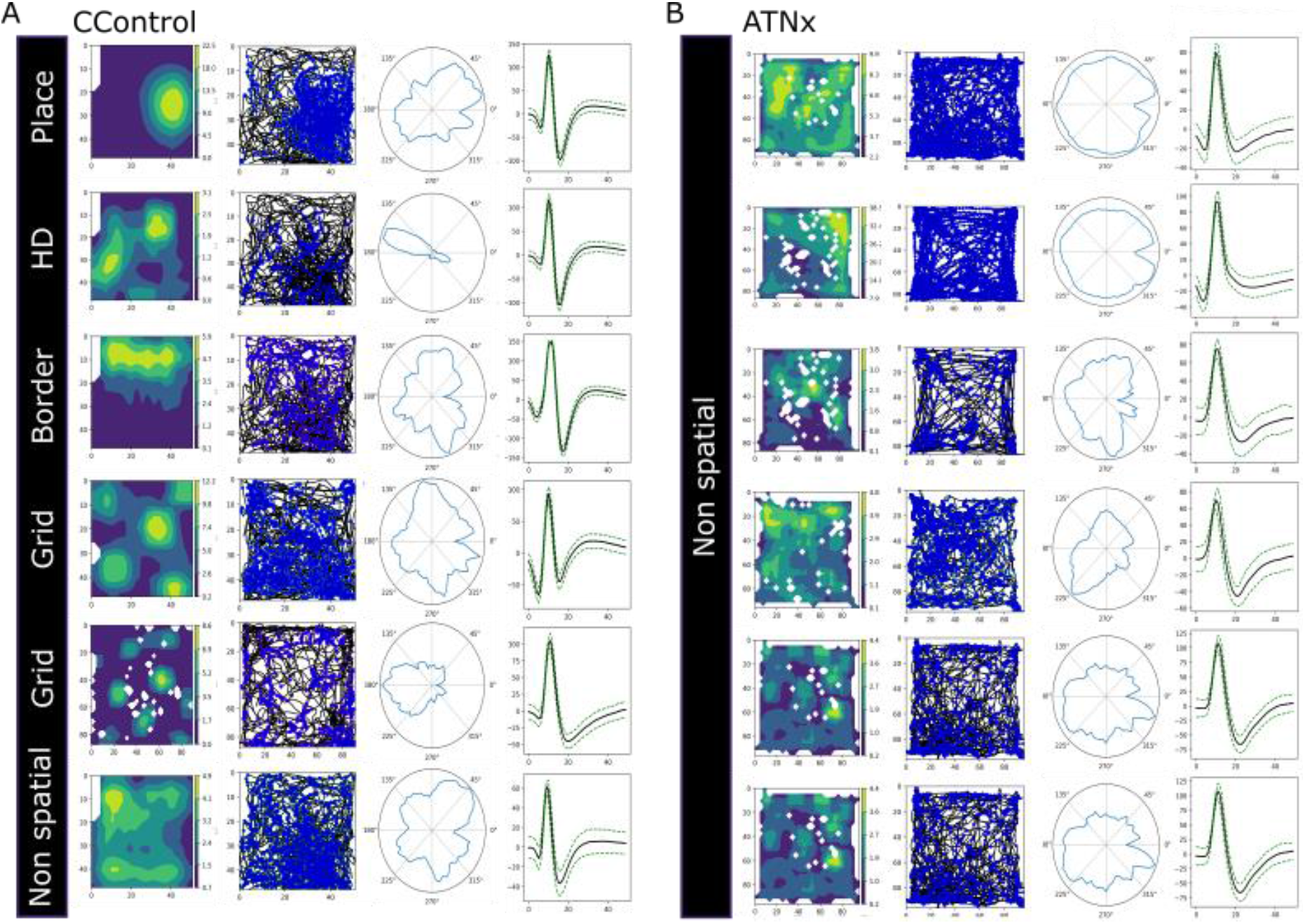
Representative single units recorded in the dorsal subiculum in control (CControl; A) and ATN lesion (ATNx; B) animals. For individual units the figures illustrate: Heatmap of spike location adjusted for time spent in each location; path in arena (black) with spike location (blue); head direction; mean spike waveform. For the CControl group, different classes of spatial cells are displayed, along with a non-spatial cell. No spatial cells were recorded in the ATNx cases. HD, head direction.

In order to assess whether this absence of spatial cells in ATNx animals was statistically significant, we first selected recordings from CControl and ATNx animals that were deemed to be independent, i.e. performed before and after electrode position was altered (during the habituation period), or were performed several days apart, as electrodes had likely shifted naturally. Recordings were considered to contain spatial cells or non-spatial cells, regardless of the number of cells recorded or whether the same cell had been recorded previously. The presence of spatial cells and non-spatial cells in the sampled recordings was accumulated in a contingency table (Table 1). The null hypothesis is that the true difference in proportion between the sample estimates is equal to 0. In other words, the percentage of recordings with cells of a certain type is independent of the condition being control or lesion. The results from running a two-sided Barnard’s unconditional exact test^46^ for a binomial model on a 2×2 contingency table consisting of the observed frequencies from the CControl and ATNx lesion recordings are as follows: for the spatial cells p < 0.0001, indicating the data strongly supports rejecting the null hypothesis; for the non-spatial cells p = 0.983, indicating that the null hypothesis should not be rejected.

**Table 1:** Recordings from CControl and ATNx animals (where a ‘recording’ is defined as a single trial of an open field recording session) were selected such that recordings were independent. Recordings were deemed to be independent if they were performed before and after adjustment of electrode position, or if they were several days apart, as the electrodes had likely shifted naturally. For each of these recordings, the presence or absence of both spatial cells and non-spatial cells were marked, and the observed frequencies are presented in the table.

| Observed frequencies for recordings | Control | Lesion | Total |
| --- | --- | --- | --- |
| **Spatial cells** |  |  |  |
| At least one spatial cell was recorded | 15 | 0 | 15 |
| Spatial cells were not recorded | 38 | 47 | 85 |
| Total recordings performed | 53 | 47 | 100 |
| **Non-Spatial cells** |  |  |  |
| At least one non-spatial cell was recorded | 16 | 13 | 29 |
| Non-spatial cells were not recorded | 37 | 34 | 61 |
| Total recordings performed | 53 | 47 | 100 |

We next conducted a Bayesian analysis on the contingency table data^47^. We find that (at about 99.99% chance), it is more likely to record a spatial cell in a CControl recording, compared to an ATNx lesion recording, P(PC > PL | D) = 0.9999. Similarly, we find that (at about 93.2% chance), it is five times more likely that we can record a spatial cell in a CControl recording compared to an ATNx lesion recording, P(PC > 5 * PL) = 0.9328. Overall, we find spatial signalling is lost in the subiculum after ATN lesions (see Fig. 4).

**Fig. 4:**
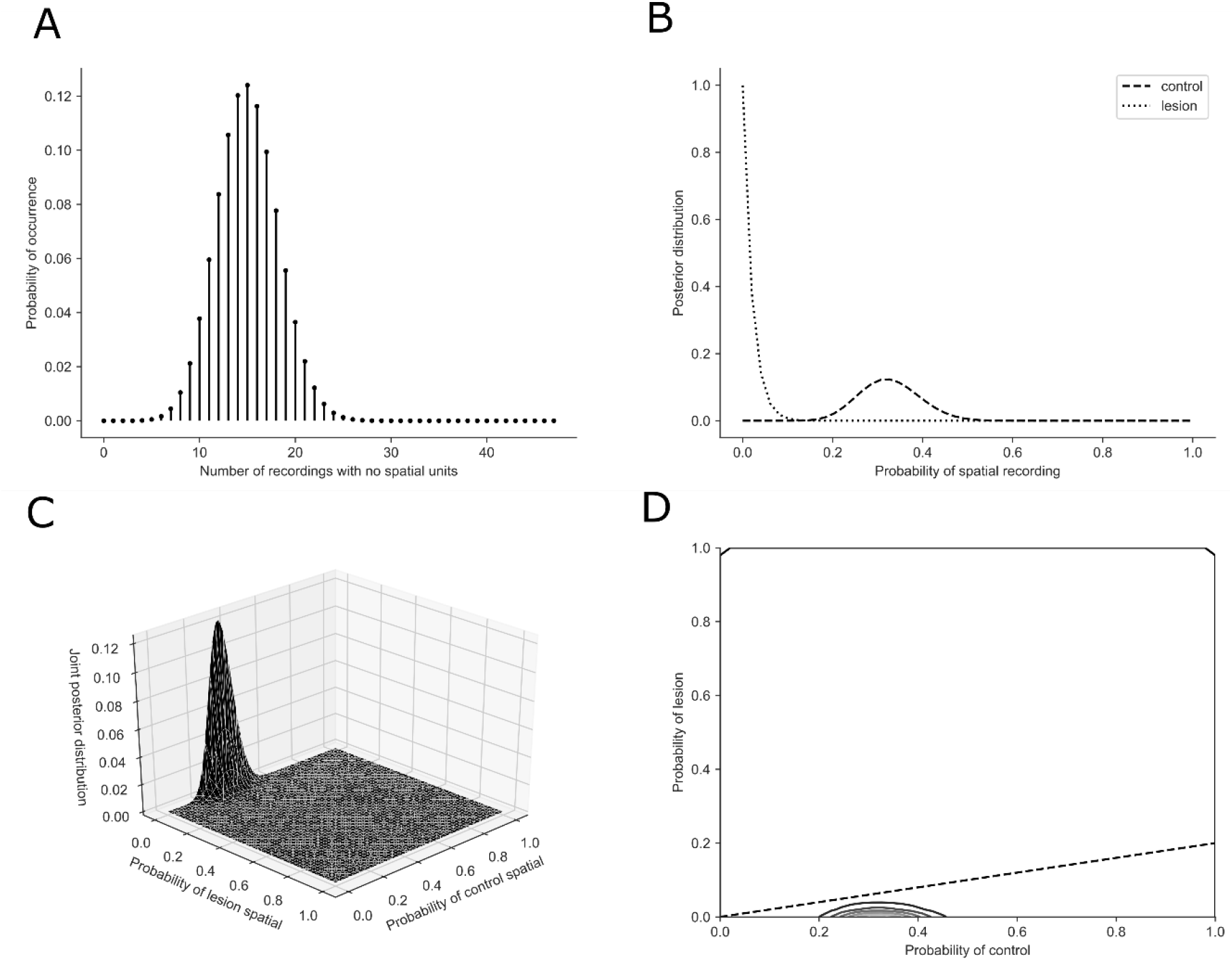
(A) The probability mass function of the random variable representing the number of recordings with a spatial cell with a number of draws equal to the number of samples from the lesion recordings, assuming that the proportion of spatial cells in the control data represents the population. (B) The posterior distribution (non-normalised) of the two spatial cell recording proportions. (C) Surface plot of the joint posterior distribution (non-normalised) of the two spatial cell recording proportions. (D) Contour plot of the joint posterior distribution (non-normalised) of the two spatial cell recording proportions, the dotted-line indicates the upper bound of the region to doubly integrate over to calculate the probability that a control recording with a spatial cell is five times more likely than a lesion recording with a spatial cell.

### ATN lesions (NMDA) leave non-spatial subiculum firing properties largely unchanged

#### Spike properties

The mean spike width of all recorded subicular cells pooled into a single group was greater in ATNx animals than CControls (CControl 155.26 ± 50.43, ATNx 213.88 ± 53.87; t = −4.14, df = 29.25, p < 0.001). Other spike properties (top section, Table 2) showed no differences between groups, including number of spikes (CControl 2694 ± 3250, ATNx 4718 ± 6326; t = −1.39, df = 22.66, p = 0.179); spike frequency (CControl 2.93 ± 4.31 Hz, ATNx 5.42 ± 7.07 Hz; t = −1.51, df = 23.79, p = 0.145); amplitude (CControl 107 ± 31 μV, ATNx 104 ± 19 μV; t = 0.47, df = 49.3, p = 0.642); height (CControl 147 ± 54 μV, ATNx 161 ± 39 μV; t = −1.34, df = 40.74, p = 0.187); and inter-spike interval (CControl 1009 ± 1381 ms, ATNx 2253 ± 3453 ms; t = −1.58, df = 21.60, p = 0.128).

**Table 2:** Summary of spike and burst properties of subicular units. Duty cycle describes the portion of the inter-burst interval during which a burst fires. * p < 0.05, ** p < 0.01, *** p < 0.001 (Welch’s Two Sample *t*-test).

|  | Control |  | Lesion |  |
| --- | --- | --- | --- | --- |
|  | Mean | ± SD | Mean | ± SD |
| Number of spikes (per 20 minutes) | 2694.93 | 3249.93 | 4718.00 | 6326.09 |
| Spiking frequency (Hz) | 2.93 | 4.31 | 5.42 | 7.07 |
| Spike amplitude (μV) | 106.91 | 31.15 | 104.33 | 19.20 |
| Spike height (μV) | 146.91 | 53.96 | 161.18 | 39.32 |
| Spike width (ms) *** | 155.26 | 50.43 | 213.88 | 53.87 |
| Inter-spike interval (ISI; ms) | 1008.59 | 1381.03 | 2253.24 | 3453.02 |
| Total bursts (per 20 minutes) | 238.90 | 435.98 | 116.6 | 157.78 |
| Total bursting spikes (per 20 minutes) | 526.59 | 981.86 | 242.90 | 331.59 |
| Mean bursting ISI (ms) | 4.08 | 0.32 | 3.88 | 0.28 |
| Spikes per burst * | 2.13 | 0.11 | 2.05 | 0.05 |
| Mean burst duration (ms) * | 5.61 | 0.64 | 5.07 | 0.45 |
| Inter-burst interval (ms) * | 11144.49 | 11073.79 | 31830.77 | 39508.75 |
| Mean duty cycle (burst duration / inter-burst interval) | 0.05 | 0.05 | 0.02 | 0.02 |
| Propensity to burst (bursting spikes / total spikes) *** | 0.12 | 0.09 | 0.06 | 0.03 |

When non-spatial cells in CControl animals were compared to putative ‘non-spatial’ cells in ATNx animals, spike width (CControl 173 ± 60 μV, ATNx 213 ± 54 μV; t = −2.56, df = 45.60, p = 0.014, Fig. 5K), and spike height (CControl 133 ± 40 μV, ATNx 161 ± 39 μV; t = −2.57, df = 42.11, p = 0.014, Fig. 5J) were both smaller in CControl animals.

**Fig. 5:**
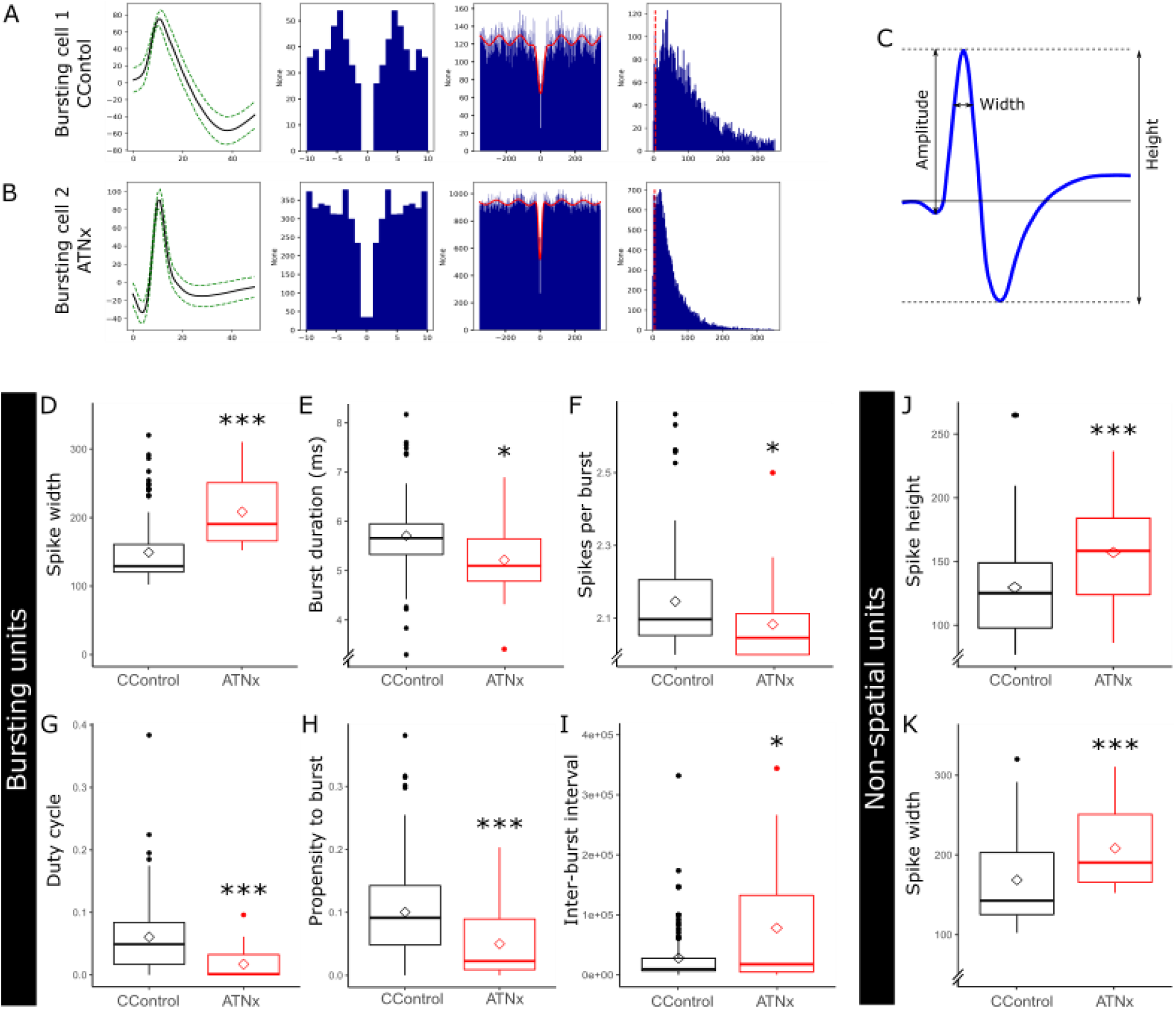
Properties of bursting and non-spatial subicular cells. (A, B) Waveforms and autocorrelation histograms were used for cell classification. (C) Diagram of waveform properties. Bursting cells in CControl showed a higher burst duration (E), had more spikes per burst (F), and had a higher propensity to burst (H). Bursting cells in ATNx had a greater spike width (D) and higher inter-burst interval (I), than non-bursting cells. (J, K) Non-spatial cells in ATN had higher spike width and spike height than non-spatial cells in CControl animals. For boxplots (D-K), filled circles indicate outliers and unfilled diamonds indicate the mean. * p<0.05, ** p<0.01, *** p<0.001 (Welch’s Two Sample *t*-test).

#### Spike properties of bursting cells

As no spatial units were recorded in ATNx animals, it was unclear whether spatial cells were present but inhibited (and, therefore, not recorded) or whether the units that were recorded were latent spatial units that were now not responding to ‘spatial’ inputs. Due to the difficulties in comparing non-spatial units in the CControls to unknown spatial or non-spatial units in ATNx animals, subicular cells were classified according to their spike properties into bursting, fast spiking, and theta-entrained cells^48^. While the percentage of subiculum bursting cells in the CControl (53% of 82 units) and ATNx (57% of 21) animals was essentially equivalent, other properties differed (lower section Table 2, Fig. 5). Cells in CControl animals showed a greater propensity to burst than those in lesion animals (CControl 0.12 ± 0.09, ATNx 0.06 ± 0.03; t = 3.69, df = 35.46, p < 0.001; Table 2). Bursting cells in CControl animals showed more spikes per burst (CControl 2.13 ± 0.11, ATNx 2.05 ± 0.05; t = 3.32, df = 31.26, p = 0.002) and greater burst duration (CControl 5.61 ± 0.64 ms, ATNx 5.07 ± 0.45 ms; t = 3.04, df = 18.21, p = 0.007), and conversely lesion animals showed larger inter-burst intervals than controls (CControl 11144 ± 11074 ms, ATNx 31831 ± 39509 ms; t = −1.56, df = 9.32, p = 0.038; Table 2).

### Muscimol infusion reversibly reduces spatial alternation memory performance to chance

When the ATN was temporarily inactivated with muscimol, spatial alternation percentage dropped to chance levels (before muscimol infusion 87.27 ± 7.41; after muscimol infusion 50.00 ± 13.97; Fig. 6B). Two animals also received bilateral saline infusions as a control, showing no deficit in spatial alternation (87.5 ± 0%). The muscimol inactivation caused a significant deficit in spatial alternation compared to both before muscimol and saline infusion (ANOVA p < 0.001, Tukey post-hoc p < 0.001).

**Fig. 6:**
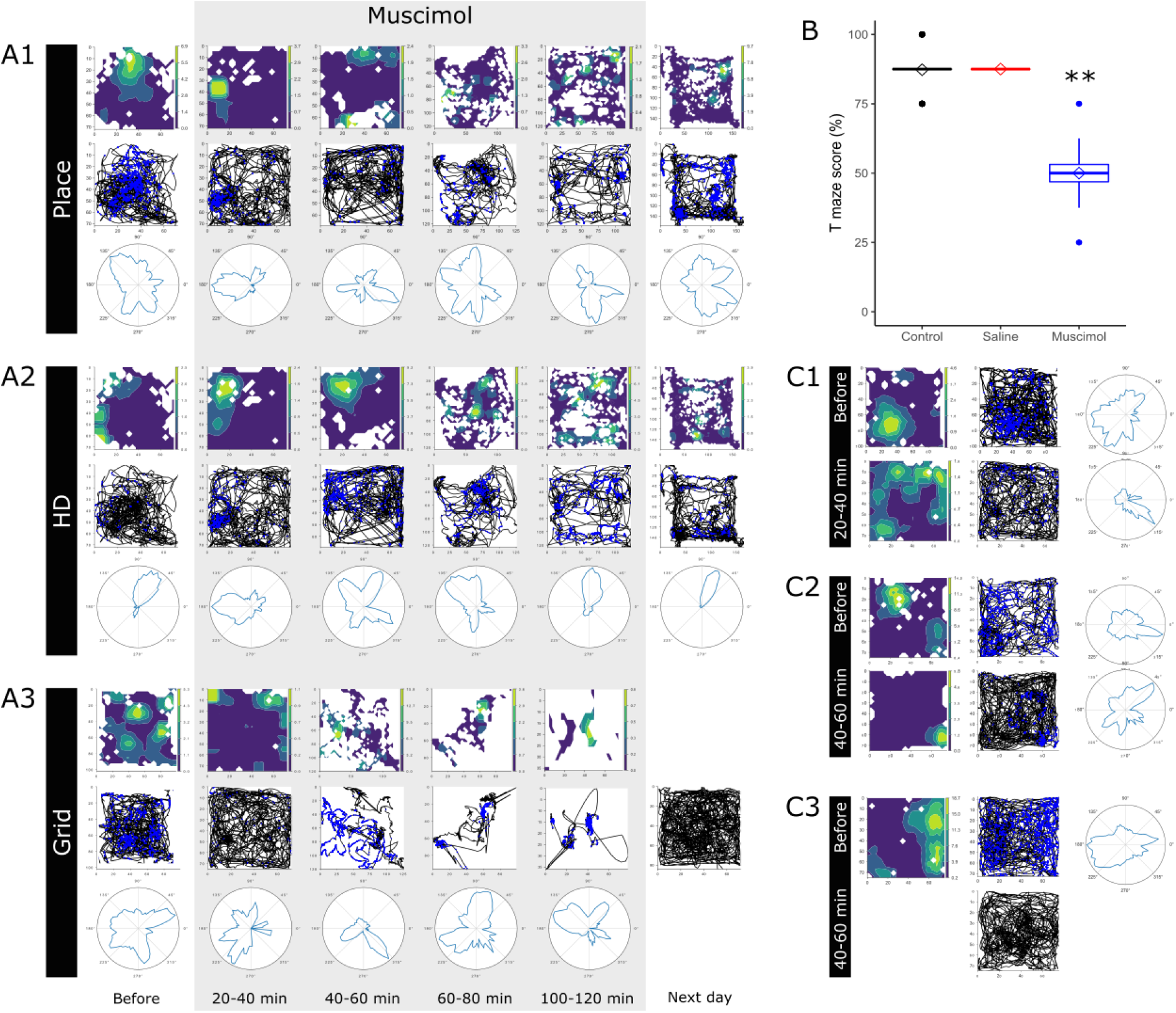
(A1-A3) Examples of spatial units before, during and after muscimol infusion. Spatial properties of single units decreased when ATN was inactivated. (Note, unit A3 showed relative inactivity after 100-120 minutes, and the cell was not recorded the next day). (B) Spatial alternation dropped to chance levels when the ATN were temporarily inactivated with muscimol, compared to the same animals prior to infusion. When saline was infused in place of muscimol no deficit was present. (C1-C3) further examples of spatial units before and after ATN inactivation. C1 shows disruption of place field shortly after muscimol infusion with some head directionality remaining, which was later disrupted. For (C3), no firing was detected after muscimol infusion. ** = p<0.01 (ANOVA with Tukey post-hoc).

### Muscimol infusion reversibly abrogates subiculum spatial firing

Prior to the muscimol infusion sessions, electrophysiological recordings were conducted on cannulated rats to allow for electrode adjustment and habituation to recording equipment. Single units were recorded during these habituation periods, however, only units recorded during the inactivation experiments were considered. Thirty-five cells were recorded during muscimol experiments. Of these, 29 were recorded at baseline, i.e., immediately prior to muscimol infusion, and so included in the study (Fig. 6 A1-A3; C1-C3).

### Muscimol infusion leaves non-spatial subiculum firing properties largely unchanged

#### Spike properties

For analysis, unless otherwise stated, Welch’s Two Sample *t*-tests were performed to consider cell parameters with respect to baseline data, with Bonferroni post-hoc correction to account for multiple comparisons of time points to baseline. When all cells were grouped together, there was no significant change in mean spiking frequency following ATN inactivation (baseline 5.07 ± 9.10 Hz, 40-60 min post infusion 2.90 ± 3.84 Hz; p > 0.20), or the number of spikes per recording (baseline 5306 ± 10344, 40-60 min post infusion 3483 ± 4607; p > 0.20). Mean spike width (baseline 180.32 ± 55.20 μV, 40-60 min post infusion 185.81 ± 50.71 μV; p > 0.20) and amplitude (baseline 113.01 ± 41.43 μV, 40-60 min post infusion 115.22 ± 45.48 μV; p > 0.20) showed no change following ATN inactivation when all cells were grouped. Finally, when all units were grouped, temporary inactivation of ATN led to an apparent doubling of the inter spike interval (ISI; baseline 664.56 ± 736.57 ms, 40-60 min post infusion 1443.29 ± 1825.34 ms) although this was not significant (p = 0.053).

Following muscimol infusion, spatial properties of subiculum cells declined, despite no decrease in firing frequency. For designated place cells prior to infusion, the place field became disrupted after ATN inactivation (Fig. 6A1). Head directionality also became disrupted without ATN input (Fig. 6A2) and grid cells did not fire in a grid-like pattern (Fig. 6A3). In the majority of cases, these spatial properties were recovered by the following day, although in some cases the cells could no longer be recorded (Fig. 6C3). Interestingly, the subiculum grid cells appeared to lose their place field initially but retained some head directionality, before this too was disrupted (Fig. 6A3, 5C1-2). We performed an additional control measure by spatially down-sampling the firing maps before and after muscimol injection to match occupancy in each spatial bin, see Supplementary Fig. 2. Statistical analysis of spatial cells is shown in Supplementary Table 1.

#### Spike properties of subiculum bursting cells

Of the 29 units included in the study, at baseline 20 were classified as bursting, 6 fast spiking, and 2 theta modulated (prior to muscimol infusion). For spike property analysis, theta modulated units were excluded due to their high firing frequency and insufficient numbers to perform further statistical analysis. When fast spiking and bursting units were combined, there was no difference in firing frequency compared to baseline, following ATN inactivation with muscimol (5.07 ± 9.10 Hz, 0-20 min post infusion 3.20 ± 4.75 Hz; p > 0.20; Fig. 8A). All animals travelled less distance, i.e. showed less activity, as time increased across the experiment (baseline 115.97 ± 33.12 m; 60 - 80 mins 58.17 ± 37.41 m; Fig. 8B), although there was no correlation between firing frequency and distance travelled (r^2^ = 0.011; Fig. 8C). Figure 8D illustrates the firing frequency of each cell at baseline, then its frequency in 5 min intervals following muscimol infusion and again the following day, in those instances where the cell was recorded again.

**Fig 7:**
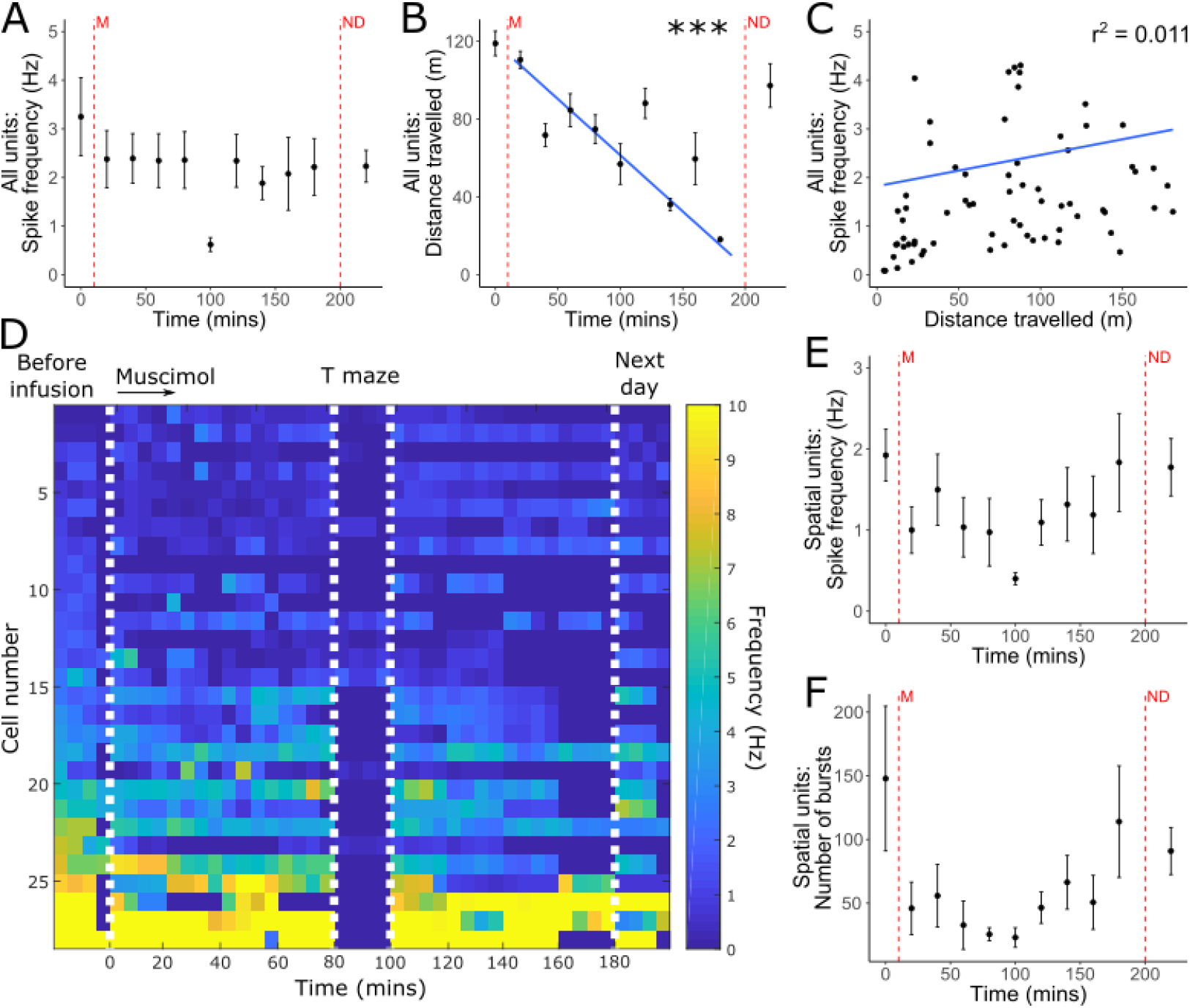
Spike properties following temporary inactivation of ATN with muscimol (M). (A) Inactivation of ATN caused no significant decrease in single unit firing frequency. (B) Animals showed decreasing levels of activity throughout the experiment, although there was no significant correlation between distance travelled and spike frequency (C). (D) Represents firing frequency of each cell recorded at baseline (left), in 5 minute bins throughout the experiment. The first white line indicates ATN inactivation with muscimol, after 15-20 minutes of baseline recording before infusion. In most cases, electrophysiological recording was paused for T maze between 80-100 minutes (second and third white lines), then continued. Recordings in which the animal was largely inactive or asleep were excluded. The final white line indicates data from the day after infusion. (E) There were no significant changes in spike firing in spatial units as a result of ATN inactivation and burst properties remained consistent including the number of bursts (F). (A, B, E, F) First red vertical line (‘M’) indicates the infusion of muscimol, second red vertical line indicates recordings taken the next day (‘ND’). Each bin represents 20 minutes of recording. Data are compared to baseline, immediately prior to inactivation, with error bars indicating SEM. Theta entrained cells are removed from A, B and C due to high firing frequency compared to other cell classes. *** p<0.001 (Welch’s Two Sample *t*-test with Bonferroni correction).

**Fig 8:**
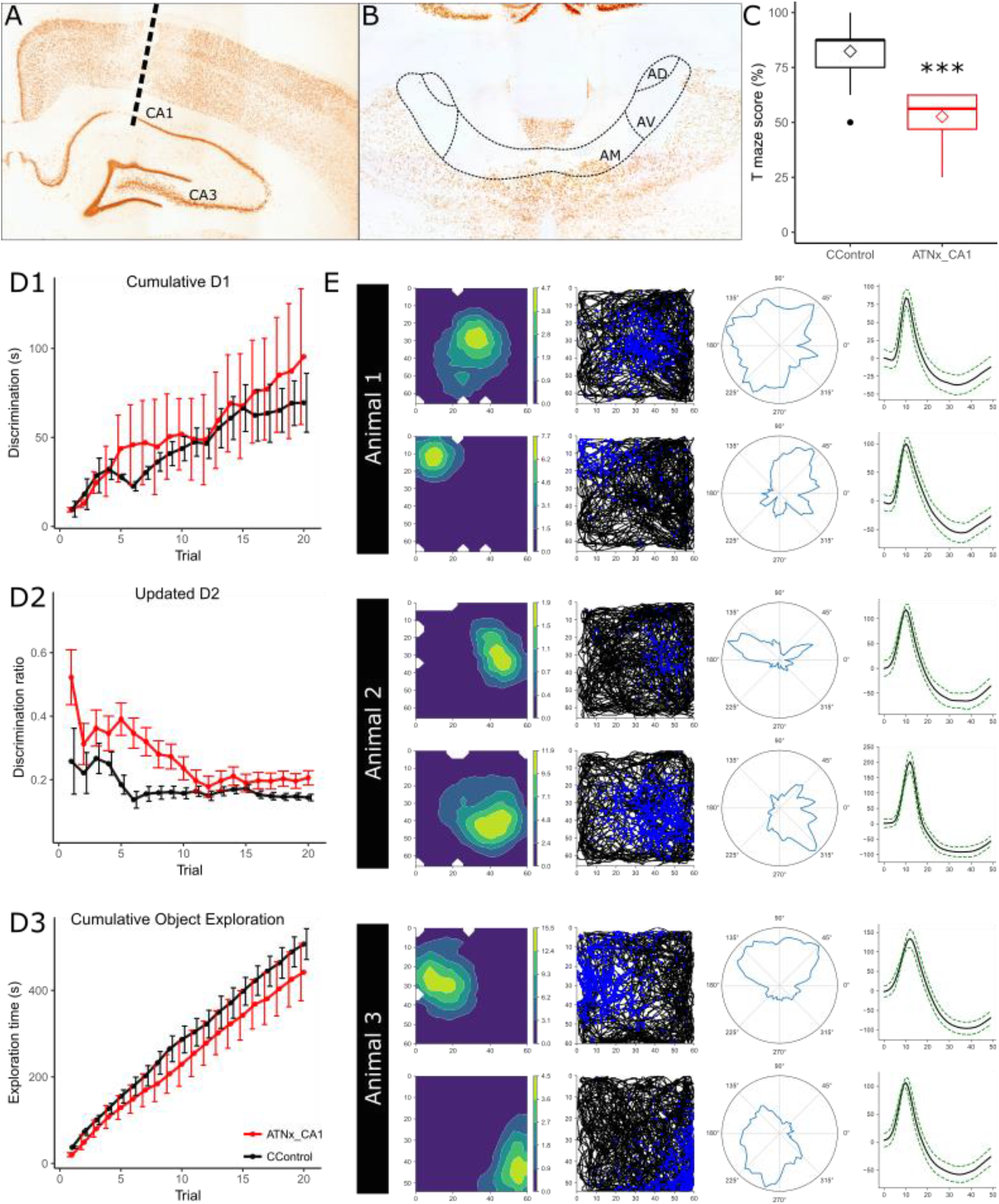
Representative electrode placement in CA1 (A) and ATN lesion (B), in NeuN-reacted sections. (C) Animals with ATN lesions and electrodes implanted in CA1 showed a significant deficit in spatial alternation task compared to controls animals (control data repeated from experiment 1). (D1-D3) The same cohort of ATNx animals showed no deficit in object recognition on bow tie maze. (E) Representative place cells recorded from CA1 in three ATNx animals.

Bursting units did not show any changes in burst properties following inactivation of the ATN, including number of bursts, spikes per burst, and burst duration (p > 0.05). Cells that had been designated as spatial units prior to infusion showed no changes in burst properties or firing frequency, indicating that these spatial cells continue to fire following ATN inactivation but no longer displayed spatial properties.

### Anterior thalamic lesions (NMDA) do not affect CA1 place cells, but reduce spatial alternation performance to chance

To explore the effects of anterior thalamic lesions on spatial processing in the dorsal hippocampus, a further three rats were implanted with a microdrive apparatus with 8 recording tetrodes into the dorsal CA1 and received bilateral, permanent (NMDA) lesions of the anterior thalamic nuclei (ATNx_CA1). As in the previous experiment, ATNx_CA1 lesions were quantified by comparing anti-NeuN reacted cell counts in the anterior thalamic nuclei with those from the CControls. In all ATNx_CA1 animals, the surgery consistently produced a marked cell loss throughout almost the entire anterior thalamic nuclei. An independent sample *t*-test showed significant difference in anti-NeuN cell counts between ATNx_CA1 (4166 ± 1003) and CControl (16209 ± 2507; p < 0.001, t = 9.08, df = 7, Welch’s Two Sample *t*-test) groups.

#### Spatial alternation deficits

As expected, ATNx_CA1 rats showed a deficit in spatial alternation on the T-maze, equivalent to ATNx animals (CControl 82.38 ± 11.18, ATNx_CA1 52.60 ± 11.40%, t = −10.71, df = 40.93, p < 0.001; Fig. 8C), confirming an impairment in spatial working memory. These animals also showed no deficit in object recognition and no difference in overall exploration time, compared to CControls (p > 0.05; Fig.8D1-3).

### CA1 place cells are unaffected by ATN lesions

Daily recordings were conducted in CA1 of ATNx animals (ATNx_CA1) performing a pellet-chasing task in an open field. From these animals, 203 well-isolated units were recorded from dorsal CA1. Units were further classified into 107 spatial units using sparsity and coherence criteria^49^ (rat 1 n = 12; rat 2 n = 67; rat 3 n = 28). Putative interneurons were considered to have been recorded from dorsal CA1 if they were recorded on the same tetrode and in the same recording session as a spatial unit. Despite the absence of spatial signals in the subiculum, hippocampal (CA1) place cells appeared intact (Fig. 8E). Properties of CA1 units are provided in Supplementary Table 2.

## Discussion

We examined the role of the anterior thalamic nuclei in the extended hippocampal system^33^, by combining *in vivo* neurophysiological recordings, permanent lesions, and temporary inactivations, with behavioural analysis. There is normal spatial firing subiculum (place, head-direction, grid, and border cells). After ATN lesions, CA1 place activity appeared preserved, whereas the usual heterogeneous subicular spatial activity was lost. Moreover, post-ATN lesion performance was at chance for spatial alternation, but intact for the object recognition memory task, indicating a specific rather than generalised deficit. The same spatial effects were reversible when using a transient or temporary inactivation: consequently, spatial firing and spatial alternation performance were reinstated after the temporary cessation of ATN activity. Thus, we conclude that anterior thalamic inputs are necessary for spatial coding in the dorsal subiculum, which contributes to spatial alternation memory performance^10^.

Lesion of the anterodorsal thalamic nucleus disrupt the parahippocampal head-direction signal but leave CA1 place cells largely intact (despite some loss of coherence and information content)^17,40^. Our more extensive ATN lesions largely left CA1 place cells intact (note, some microstructural changes, alterations in neuronal activity, and synaptic integrity changes have been reported in CA1 following ATN disruption)^40, 52^. Notably, the proportions of place cells in CA1 found in ATNx-CA1 animals is in line with our own previous experience^52, 53^, as well that of others^54,55,56,57^ and stands in contrast to the more disruptive effects of medial entorhinal cortex lesions on CA1 place cell activity^58^. Given that the majority of pyramidal cells in CA1 are place cells and that CA1 heavily innervates the subiculum, it could be assumed that many subiculum spatial cells are principally driven by their CA1 inputs. The present study revealed, however, that CA1 projections alone do not support subicular firing or spatial alternation memory. Projections from the ATN, presumably both direct and indirect, are also crucial for subicular spatial cellular discharge, alongside (behavioural) spatial alternation. In addition to the head-direction cells in the anterodorsal thalamic nucleus, the anteroventral nucleus possesses theta-modulated head-direction cells^59^, while the anteromedial nucleus contains place cells and perimeter/boundary cells^16,60^, suggesting that ‘early’ spatial signals are present in ATN. These results reaffirm the view that the ATN have critical roles in navigation and spatial learning, and spotlight the pivotal role of the subiculum, given the severity of the alternation deficit, despite the sparing of CA1 place cells. As the subiculum is a principal source of hippocampal projections beyond the temporal lobe, these findings also reveal how anterior thalamic damage might indirectly impact upon a wide array of other sites, including the mammillary bodies and retrosplenial cortex (RSC)^61,62^.

Consistent with previous reports^33–36^ we found spatial cells, including grid-like cells, in the dorsal subiculum of healthy, control animals; however, no spatial units were recorded in the dorsal subiculum of ATN-lesioned animals. Within-animal trials were used to confirm this phenomenon, whereby the ATN was temporarily inactivated using micro-infusions of muscimol. Spatial activity was lost as muscimol took effect, and recovered as muscimol was metabolised. These effects, which presumably arise from the loss of AM and AV discharge to the subiculum, alongside AD disconnection from the pre-, post-and parasubiculum, show the closeness of this relationship.

Given the routes of ATN fibres reaching the medial temporal lobe, it is worthwhile considering whether we might have recorded from fibres of passage. However, even *in vitro*, “direct recording of single AP (action potential) transmission is challenging” due to the small diameters of axons and recording instability^63^, whereas we have recorded for long durations in freely-behaving, implanted animals. Furthermore, our units have a half-width of ~0.8-1.2msec: precisely within the refractory period for action potentials. A further issue concerns potential disruption to thalamic fibres of passage, despite our use of cytotoxic lesions. The finding that reversible ATN lesions with muscimol, which do not affect fibres of passage^64^, disrupted grid cells (and all other spatial subicular cell classes) confirms that the surgeries targeted ATN neurons.

Although grid cells have been widely studied in the hippocampal formation since their discovery in the medial entorhinal cortex (mEC)^55^, the presence of grid cells in the dorsal subiculum is still a relatively new finding^36^. Grid cells had, however, been previously reported in the pre- and parasubiculum^65^; parahippocampal areas that receive inputs from the anterior thalamic nuclei^66^. Grid cell (and head-direction) activity in these same parahippocampal areas is disrupted by anterior thalamic lesions^17^. The grid-like signal found in presubiculum and subiculum might, therefore, be partly thalamic^17^, as well as entorhinal, in origin.

Stewart and Wong^67^ originally observed *in vitro* that subicular cells can be classified into bursting and non-bursting classes (confirmed by Sharp and Greene^68^ *in vivo*). Subsequently, Anderson and O’Mara^48^, in a fuller *in vivo* analysis, concluded subicular units could be separated into bursting, regular spiking, theta-modulated, and fast spiking units; furthermore, Simonnet and Brecht find sparsely bursting subiculum cells may carry more spatial information^69^. Here, we found that although the proportions of bursting cells seem unchanged, their properties were altered after anterior thalamic lesions. The proportion of bursting cells in the CControl and ATNx groups remained equivalent, but some of their other phenotypic properties differed: controls showed a greater burst propensity, as well as having more spikes per burst and greater burst duration. Conversely, lesion animals showed somewhat larger inter-burst intervals than controls. This alteration in bursting may affect the fidelity of subicular-retrosplenial transmission, as bursting cells in dorsal subiculum with direct connections to granular RSC have a direct impact on sharp wave ripples in RSC^70^.

The behavioural effects of the ATN lesions largely matched those of hippocampal lesions: there was severely impaired T-maze alternation, but spared object recognition memory in the bow-tie maze^71^. We also found somewhat increased motor activity in ATNx rats in square arenas, matching prior evidence of activity increases following ATN lesions in spatial settings^7,72,73^ potentially linked to a failure to encode and retain locations, a finding also seen after hippocampal damage^74,75^.

The subiculum is the primary hippocampal output of area CA1^41,42,32,76,77^, an area with a substantial preponderance of place cells, thus ensuring that the CA1 input to the subiculum is principally spatial. Nevertheless, our data cast new light on this relationship as CA1 inputs are not sufficient to ensure subiculum place cell firing. One possibility is the ATN normally exert a direct modulatory, including oscillatory^78^, influence on the subiculum that, when removed, leads to changes in gain control that disrupt information processing. Given the spatial cells in the ATN^14–16^ such an input might, for example, normally help hippocampal and parahippocampal regions co-register their various spatial signals, including those from CA1. Such a role might help explain that while the ATN lesions increased dorsal subiculum spike width and reduced bursting properties, they left many firing features seemingly intact, e.g., overall spike frequency and amplitude. Likewise, immediate-early gene analyses have found that although ATN lesions cause hypoactivity in parahippocampal fields, but may have little gross impact on the dorsal subiculum^51,74^. One shortcoming with this account is that it fails to explain why the direct CA1 projections to entorhinal and prefrontal areas (which do not require the subiculum) are not sufficient to protect against disruptions to spatial learning and memory.

Such considerations invoke a much wider, network impact of ATN loss. This impact includes direct efferent actions on the subiculum, alongside indirect actions via parahippocampal and retrosplenial targets. As noted above, there remains the issue of why other areas (e.g., CA1, entorhinal cortex) are not sufficiently independent of the ATN to preserve subicular spatial firing and effective spatial memory. It may, therefore, be relevant that the subiculum has very dense direct and indirect projections to the ATN that are part of complex, reciprocal set of pathways^33^. Furthermore, the direct interactions between these two sites are needed for T-maze alternation, irrespective of whether they are ATN projections to the dorsal hippocampus (including subiculum) or subiculum projections to the ATN^10^. Extending our understanding these reciprocal actions will prove integral to appreciating why ATN damage is such a critical component of diencephalic amnesia^5^.

Overall, the data here show that ATN inputs are necessary for spatial activity in the subiculum and contribute to spatial alternation memory performance. Further, these data indicate that signalling within the rest of the hippocampus (e.g., CA1) is not sufficient for the varied spatial signals found in the subiculum. We suggest that subicular spatial signals arise from converging inputs from CA1, the various anterior thalamic nuclei, and parahippocampal areas, including entorhinal cortex. Further work is required to determine the nature of the computations performed by the subiculum on the inputs it receives from ATN; how transformation rules are applied to these inputs to support spatial alternation performance; and to discover by what means the substantial input from hippocampal area CA1 is gated by convergent ATN projections reaching the subiculum.

## Acknowledgements

This work was supported by a Joint Senior Investigator Award made by The Wellcome Trust to JPA and SMOM (103722/Z14/Z).

## Author contributions

BEF: Acquisition of data, analysis and interpretation of data, drafting or revising the article;

SKM: Developing analysis algorithms, Python script writing and validation, analysis and interpretation of data, drafting or revising the article

MC: Acquisition of data, analysis and interpretation of data

MNI: Original conception and development of NeuroChaT

JPA, SMOM: Conception and design, analysis and interpretation of data, drafting and revising the article

## Table of Methods

### Key Resources Table

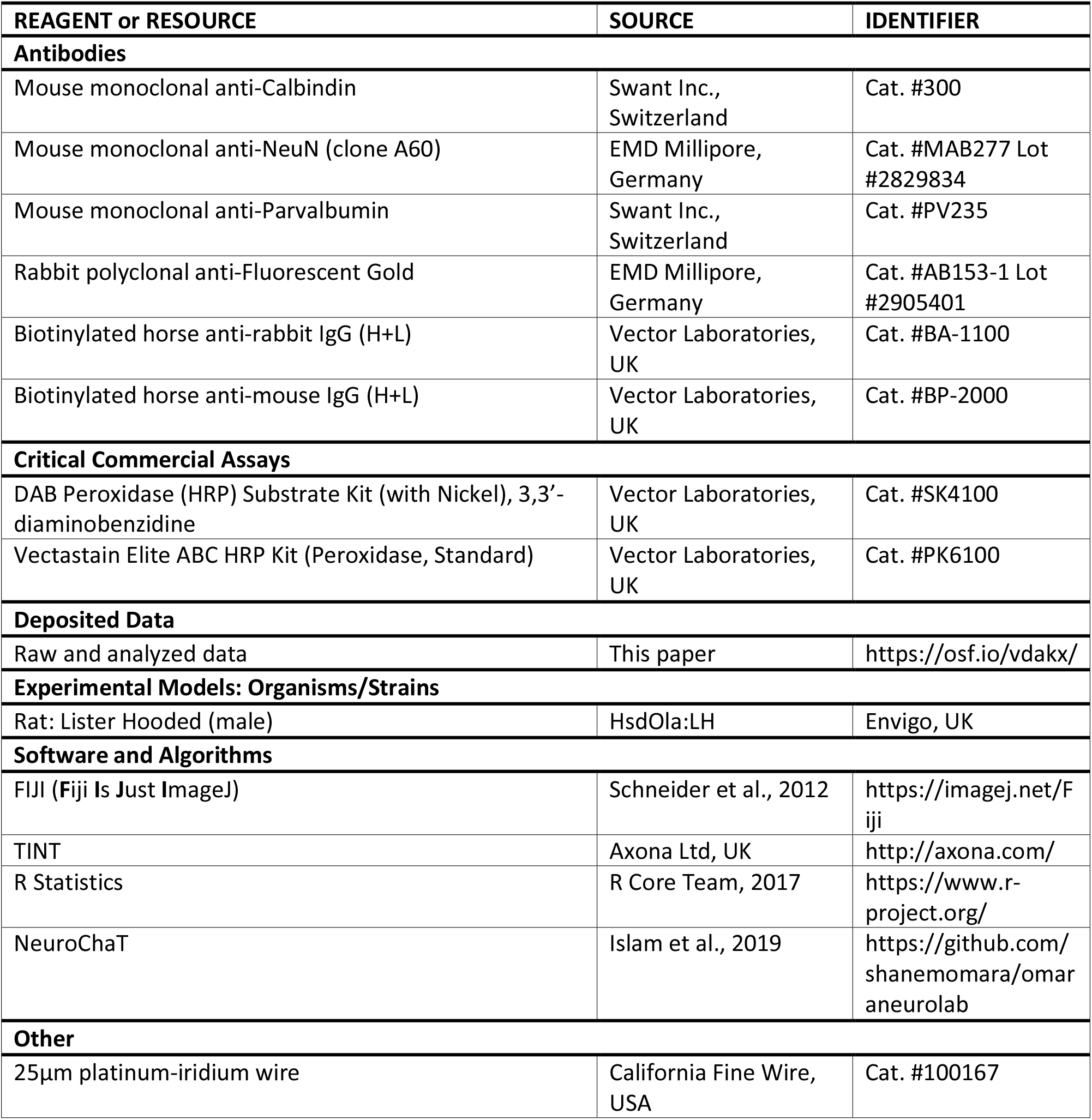

### Lead Contact and Materials Availability

Requests for further information and resources should be directed to and will be fulfilled by the Lead Contact, Shane O’Mara. This study did not generate new unique reagents.

### Experimental Model and Subject Details

Experiments were conducted on 23 male Lister Hooded rats (Envigo, UK) with pre-procedural weights of 309-356g. Upon arrival, animals were cohoused on a 12-hour day/night cycle and handled daily by the experimenter for a week before surgical procedure. Prior to surgery and during recovery, animals had free access to food and water; during behavioural testing, food was restricted, but ensured the animals did not fall below 85% of the animal’s free feeding weight. All rats were naïve prior to the present study. Selection of animals between lesion and control groups was alternated according to body weight prior to surgery (starting with the heaviest), so that pre-procedural weights were matched and the groups balanced.

## Method Details

During stereotaxic surgery, rats were implanted unilaterally with twenty-eight electrodes of 25μm thickness platinum-iridium wires (California Fine Wire, USA) arranged in a tetrode formation. Tetrodes were targeted at the dorsal subiculum, CA1, or at the subiculum and CA1 simultaneously. An additional bipolar electrode (stainless steel, 70μm thickness) targeting the ipsilateral retrosplenial cortex (RSP) was also implanted in all but two cases (see below), but the data from these electrodes are considered elsewhere. All electrodes were connected to a 32-channel microdrive (Axona Ltd., UK).

For the permanent lesion experiments, seven animals received stereotaxic cytotoxic lesions targeting the anterior thalamic nuclei (‘ATNx’), followed by electrode implantation targeting dorsal subiculum and RSP. Meanwhile, three rats underwent sham injections of equivalent volumes of PBS only (‘Sham’ controls). A further four rats (‘Normal’ controls) had no sham lesion procedure, i.e., just had electrodes implanted. Both the sham and normal controls had electrodes targeting dorsal subiculum.

The temporary inactivation (muscimol) experiment, followed the permanent lesion study. For this, a further six animals (ATNmusc) were implanted with tetrode and bipolar electrode configurations alongside bilateral infusion cannulae (26 gauge, 4mm length, Bilaney Consultants Ltd., UK) in the ATN. Electrodes were positioned as above, with two exceptions; one animal was implanted with four tetrodes targeting the subiculum, three targeting CA1 and one targeting retrosplenial cortex, with no additional bipolar electrode; and one rat was implanted with tetrodes targeting the subiculum and the bipolar electrode targeting CA1.

Finally, for the CA1 experiment, a further three rats (ATNx_CA1) received permanent bilateral ATN lesions followed by electrode implantation into CA1.

### Ethics

All experimental procedures were in accordance with the ethical, welfare, legal and other requirements of the Healthy Products Regulatory Authority regulations, and were compliant with the Health Products Regulatory Authority (Irish Medicines Board Acts, 1995 and 2006) and European Union directives on Animal Experimentation (86/609/EEC and Part 8 of the EU Regulations 2012, SI 543). All experimental procedures were approved by the Comparative Medicine/Bioresources Ethics Committee, Trinity College Dublin, Ireland prior to conduct, and were carried out in accordance with LAST Ireland and international guidelines of good practice.

### Surgical methods - permanent ATN lesions and electrode placements

Rats were first anaesthetised with isoflurane (5% to induce anaesthesia, 1-2% to maintain) combined with oxygen (2 L/minute). After being placed in a stereotaxic frame, chloramphenicol 0.5% eye gel, pre-operative antibiotics (Enrocare, 0.1ml in 0.5ml saline) and analgesia (Metacam, 0.1ml) were administered.

The skull was exposed and connective tissue removed. For the ATNx cohort (n = 7), bilateral neurotoxic lesions targeting the ATN were performed using slow infusions of 0.12M *N*-methyl-D-aspartic acid (NMDA) dissolved in phosphate buffer solution (PBS, pH 7.35). NMDA was infused over 5 minutes (0.22 or 0.24μl per site) via a 0.5μl Hamilton syringe (25 gauge), with the syringe left in position a further 5 minutes at each of four target sites before slow retraction. The craniotomies were then sealed using bone wax (SMI, St Vith, Belgium). The ATN lesion coordinates, with the skull flat, were as follows from bregma: AP −1.7mm, ML ±0.8mm, DV −5.7mm from top of cortex; AP −1.7mm, ML ±1.6mm, DV −4.9mm from top of cortex. Sham control animals (n=3 rats) underwent four equivalent infusions of PBS only.

Bundles of 28 electrodes of 25μm thickness platinum-iridium wires (California Fine Wire, USA) arranged in a tetrode formation were implanted unilaterally. Tetrodes were implanted aimed at the dorsal subiculum (AP −5.6mm, ML 2.5mm, DV −2.7mm from top of cortex), CA1 (AP −3.8mm, ML 2.5 mm, DV −1.40mm from top of cortex), or both. Electrodes were stabilised with dental cement (Simplex Rapid, Kemdent, UK) attached to the screws implanted into the skull. An additional bipolar electrode (stainless steel, 70μm thickness) targeting the ipsilateral retrosplenial cortex (RSP) was also implanted in all but two cases, the data from which are considered elsewhere. All electrodes were connected to a 32-channel microdrive (Axona Ltd., UK).

For the ATN inactivation experiment, a further six animals (ATNmusc) were implanted with tetrode and bipolar electrode configurations alongside bilateral infusion cannulae (26 gauge, 4mm length, Bilaney Consultants Ltd., UK). Cannulae were placed targeting ATN (AP −1.7 mm, ML ± 3.8 mm, DV - 4.0mm from top of cortex, at angle 28.6° towards centre), then fixed in position using dental cement and dummy cannulae inserted to prevent blockage. Electrodes were positioned as above, targeting dorsal subiculum and RSC, with two exceptions; one animal was implanted with four tetrodes targeting the subiculum, three targeting CA1 and one targeting retrosplenial cortex, with no additional bipolar electrode; and one rat was implanted with tetrodes targeting the subiculum and the bipolar electrode targeting CA1.

Glucosaline (5-10ml) was administered subcutaneously post-operatively and the animal allowed to recover. Animal weight, activity, and hydration were closely monitored daily for a minimum of 7 days before beginning electrophysiological recordings.

### Electrophysiological recordings

Electrophysiological recordings were obtained using an Axona Ltd (UK) 64-channel system, allowing dual recordings of single units and local field potentials (LFP) from each electrode. Initial habituation recordings were conducted in a 60 x 60 cm square, walled arena (height 42 cm). Later testing involved a larger arena (105 x 105 cm, 25 cm height). For both arenas, the walls and floors were made of wood painted matt black. A black curtain could be closed around the arena to remove distal spatial cues, and visual cues could be attached to the curtain as required. The habituation sessions in the small arena allowed the animal to acclimatise to the recording procedure and the experimenter to adjust electrode locations until the optimal recording depth was reached. After the habituation period (usually 3-7 days), rats were first trained on the behavioural tasks (T-maze then bow-tie maze) with up to one hour of free exploration with pellet-chasing tasks typically recorded afterwards in the same day. Pellet-chasing included the rotation of spatial cues on the curtain during the recording of single unit activity in the small arena to examine whether spatial units remap accordingly, exploration in the small then large arena in order to assess spatial cell remapping, and consecutive recordings to examine sleep properties (data not described here).

### Behavioural tasks

#### Spatial alternation

A four-arm cross-shaped wooden maze with raised sides (119 x 119 cm full length; each arm 48 x 23 cm; height 30 cm) was used for the spatial alternation task, allowing the rotation of start points. Each arm could be blocked close to the centre to form a T-maze. In addition, a barrier could be placed within an arm to form a holding area for the start position. Distal spatial cues were available in the recording room including the pulled-back curtain, electrophysiological recording equipment set on wall-mounted shelves, a desk and computer. Animals were first habituated in pairs to the maze and allowed to freely explore for 10 minutes. Rats then individually had two pre-training sessions (5 minutes each) in which they were first placed behind a barrier at the start position at one end of the maze, retained for 10 seconds, then the door removed and the rat allowed to explore the maze. The maze arm opposite to the start was blocked and sucrose pellets (TestDiet 5TUL 20mg, USA) were placed in a shallow dish at the end of each open arm so that they were not visible from the centre of the maze. Rats learnt to run to an arm of the maze to obtain sucrose pellets, which were replaced once they had been consumed and the animal left that arm. The maze was then altered so that a different start arm and blocked arm were used, and another training session run. Each rat had four 5-minute training sessions per day such that each start/blocked configuration was experienced (opposite, adjacent, opposite) for two days initially, with an additional day if required.

For the experiment, rats were placed in the start position for 10 seconds. In each trial, there was an initial forced run (‘sample’), in which two arms of the cross-shaped maze were blocked forcing the animal into either the left or right arm in order to obtain two sucrose pellets from the end of the arm. The animal was then picked up and returned to the start position and held for approximately 20 seconds before being released for the choice run (‘test’), in which one of the barriers was removed from the maze so that the animal had the choice of either the left or right arm (Fig. 2A). Sucrose pellets were only available in the arm opposite to the sample run, so the rat was rewarded if it alternated. A choice was determined when the back paws of a rat had entered the arm. The rat was then removed from the maze and placed in an open holding cage for 2-3 minutes whilst the maze was rearranged. Each animal had eight sessions consisting of eight trials each, with both the start position and forced turn pseudorandomised so that the same arrangement did not occur more than twice consecutively. Rats were connected to the electrophysiological recording equipment throughout pre-training and each test session.

#### Bow-tie maze

The bow-tie maze allows continuous object recognition testing, with multiple trials and new novel objects during each session^71^. The bow-tie shaped maze had raised sides (wood painted matt black; 120 cm long x 50 cm wide, 50 cm height) and a central sliding. Partitions at each end of the maze split both ends into two short corridors (Fig. 2C). The animals were first habituated to the maze for 10 minutes, by allowing free exploration with sucrose pellets scattered throughout. Next, during the 10 minute session, pellets were first placed in wells at the two ends of the maze and the rat trained to run from one end of the maze to the other when the central door was opened. Then, opaque plastic objects (a funnel and a beaker) were placed behind each baited well. The objects were gradually moved so that they increasingly covering the wells. Rats underwent 4-5 ten minute pre-training sessions, until they readily shuttled across the maze to retrieve sucrose pellets by pushing objects.

For the experimental procedure, 22 pairs of novel objects were used. In the first trial, one novel object was placed covering the sucrose pellets. The rat retrieved the pellets and investigated the object at that end of the maze. After 1 minute, the central door was opened and the rat passed to the opposite end of the maze, where there would be a repeat of the original object (now familiar) and a new novel object (both covering sucrose pellets). This procedure was repeated for all 22 pairs of objects, so that each of the 21 trials consisted of a new novel object and the previous object, which was now familiar^71^. Animals were video recorded throughout.

For analysis, the time spent investigating each object was recorded and two measures of recognition, D1 and D2, were calculated^71^. D1 represents the difference in exploration time between novel and familiar objects, and is calculated by subtracting the time spent exploring a familiar object from the time spent exploring a novel object. Cumulative D1 represents the sum of the exploration time for all novel objects minus the sum of exploration time for all familiar objects across all trials. D2 represents the total difference in exploration time (cumulative D1) divided by the total exploration time for both novel and familiar objects, resulting in a ratio that ranges between +/- 1.

### Transient inactivation of the ATN (muscimol)

An additional cohort (n=6, ATNmusc) was implanted with bilateral infusion guide cannulae (26 gauge, 4mm length; Bilaney Consultants Ltd., UK) aimed at the ATN at an angle of 28.6° towards the midline (AP −1.7mmg, ML ±3.6mm, DV −4.0mm from top of cortex) alongside subiculum electrode implantation. A dummy cannula (0.203mm diameter, 4mm length; Bilaney Consultants Ltd., UK) was use to protect each guide cannula during recovery and normal recording activity. All other surgical and electrophysiological methods were the same as those used for the animals with permanent ATN lesions.

Following recovery, rats were trained daily for a minimum of one week prior to commencing inactivation experiments. Rats were lightly restrained and the dummy cannulae removed and replaced several times. During this period, rats were also trained in the spatial alternation (T-maze) task and electrophysiological activity was recorded during free exploration and pellet-chasing in the large and small square arenas. All apparatus matched that used for the ATNx rats.

On the day of experimentation, electrophysiological recordings of exploration and pellet-chasing were first performed pre-infusion for 20-40 minutes to establish a baseline. The animal was then lightly restrained and muscimol (concentration 0.5mg / 1ml saline) infused through a 33gauge infusion needle with a 1- or 2mm projection past the length of the implanted guide cannula, targeting the ATN. Muscimol was infused over 90 seconds using a 0.5μl Hamilton syringe and infusion pump (KD Scientific, Hollister, USA). The infusion needle was retained in position for a further 60 seconds before it was removed and replaced with the dummy cannula. Rats received a 0.2μl at each of two locations per hemisphere using both a 5- and 6mm length infusion needle in order to target the whole ATN (Fig. 1H). Following infusion, electrophysiological recordings during pellet-chasing and exploration were conducted in consecutive 20 minute sessions in the small and large square arena. Arenas were swapped between recordings in order to maintain the animal’s interest in the surroundings and to assess whether spatial remapping occurred. Between 90 and 120 minutes into the experiment, the T-maze task was performed (8 trials, pseudo-randomised starting points). Animals were then returned to the square arena and recorded for a further 2-3 hours, including during sleep. Regular diet and water were freely available in the recording arena after the T maze test. The following day, further electrophysiological recordings were undertaken to determine whether the effects of muscimol had ceased and cell activity had returned to baseline.

Muscimol infusions were repeated 1-2 weeks after the initial experiment. In two cases, an additional control infusion of saline was given. In order to visualise the location of the muscimol infusion, the tracer fluorogold (Sigma-Aldrich Ireland Ltd, Ireland) was infused one day prior perfusion.

### Perfusion and histology

After completion of experiments, animals were sacrificed and perfused transcardially with 0.1M phosphate buffered saline (PBS) then 2.5% paraformaldehyde (PFA) in 0.1M PBS. Brains were removed and post-fixed in PFA for 24 hours then transferred to 30% sucrose in 0.1M PBS solution for 2 days. A cryostat (Leica CM1850) was used to cut 40μm sections in a 1:4 series. One series was mounted onto double gelatin-subbed microscope slides and, once completely dry, washed in decreasing concentration of alcohol (100, 90, 70%) before being stained with cresyl violet, a Nissl stain (Sigma-Aldrich Ireland Ltd, Ireland). Sections were then dehydrated with increasing alcohol concentrations, washed in xylene, and cover-slipped.

Of the other series, one was reacted against anti-calbindin antibody raised in mouse (Swant Inc., Marly, Switzerland) and another against anti-NeuN antibody raised in mouse (EMD Millipore, Germany). The remaining series was reacted with either anti-parvalbumin antibody raised in mouse (Swant Inc., Marly, Switzerland) or, in the temporary inactivation cohort, with anti-fluorogold raised in rabbit (EMD Millipore, Germany).

In brief, sections were washed in a quench solution (10% methanol and 0.3% hydrogen peroxide in distilled water) before PBS (0.1M pH 7.35) then PBST (2ml Triton X-1000 in 1 litre 0.1M PBS; pH 7.35) washes. Sections were stirred for one hour in 4% normal horse serum in PBST before the primary antibody was added (1:5000 dilution in PBST for calbindin, parvalbumin and fluorogold; 1:10000 for NeuN) and stirred at 4° overnight. Sections were then washed in PBST before being incubated for two hours in 1:250 dilution of horse-anti-mouse (Vector Laboratories, UK), or, in the case for fluorogold, horse-anti-rabbit (Vector, UK), in PBST. After further PBST washes sections were incubated at room temperature in Vectastain Elite ABC Solution (Vector Labs, UK) before further PBST and PBS washes. Sections were then reacted with DAB solution (Vector Labs, UK) and washed in PBS.

All sections were mounted on double-subbed slides, and fluorogold reacted slides were lightly stained with cresyl violet for improved tissue visualisation before coverslipping. Sections were imaged using either an Olympus BX51 upright microscope or Leica Aperio AT2 slidescanner.

## Quantification and Statistical Analysis

### Behavioural analysis

To analyse T-maze results, the mean score was compared between Normal Control, Sham Control and ATNx groups using ANOVA with Tukey post hoc test. For the muscimol experiments, T-maze results during ATN inactivation were compared to the same animals before inactivation.

To analyse object recognition using the bow-tie maze, the time spent investigating each object was recorded and two measures of recognition, D1 and D2, were calculated^71^. D1 represents the difference in exploration time between novel and familiar objects, and is calculated by subtracting the time spent exploring a familiar object from the time spent exploring a novel object. Cumulative D1 represents the sum of the exploration time for all novel objects minus the sum of exploration time for all familiar objects across all trials. D2 represents the total difference in exploration time (cumulative D1) divided by the total exploration time for both novel and familiar objects, resulting in a ratio that ranges between +/- 1.

### Unit identification and isolation

Spike sorting was performed automatically in Tint using k-means (Axona Ltd., Herts, UK) and cluster cutting refined manually. Unit identification used the following criteria: units had to be active, and show consistent waveform characteristics (amplitude, height, and duration) during recording, as well as a clean refractory period (>2 ms) in the inter-spike interval (ISI) histogram. Spike amplitude was measured as the difference between the positive peak and first negative peak before the positive peak, if present, or zero. Spike height was the difference between the spike peak to the minimum value of the spike waveform. Spike width was the distance in microseconds beyond which the waveform drops below 25% of its peak value.

Units were sometimes seemingly recorded for more than for 1 day, despite electrode lowering. For these cases, cells were monitored on the relevant tetrodes from day-to-day; for analysis, only clean recordings with the largest sample size and spikes of the highest amplitude were chosen. To avoid double-counting cells, care was taken to exclude seemingly-related samples from analysis. During spike sorting, the signals from each cell were carefully followed from first appearance to complete loss, to avoid overestimation of cell counts. Once well-defined neuronal signals were isolated recording commenced. For permanent lesion experiments, rats had to explore at least 90% of the open field in a session to be included in analyses to allow reliable calculation of spatial characteristics.

Standard statistical testing was performed using an open-source custom-written suite in Python (NeuroChaT^81^; available for download at https://github.com/shanemomara/omaraneurolab), and additional custom codes in R (R Foundation for Statistical computing, Vienna, Austria, https://www.r-project.org). Units were classified based on the spatiotemporal features of their activity in the open field during pellet-chasing, as described below.

#### Bayesian analysis

To investigate whether the apparent absence of spatial signal in ATNx animals was significant, a Bayesian approach was applied as follows. Let PC denote the probability that a control recording will contain a spatial cell, and PL denote the probability that a lesion recording will contain a spatial cell. Let D denote the recorded data. By Bayes theorem, P(PC, PL | D) = P(D | PC, PL) * P(PC, PL) / P(D) (i.e. posterior = likelihood * prior / probability of the evidence). We use a uniform prior distribution, as we have no prior belief of how common recordings in the subiculum with spatial cells are, so P(PC, PL) = 1. The probability of the evidence is a constant normalising parameter, which can be calculated be ensuring the posterior distribution sums to 1. By the independence assumption of selected recordings, the likelihood function can be modelled as the product of evaluating the probability mass functions arising from two binomial distributions:

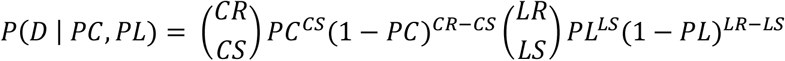

Where the data D, provides the information; CR: the number of control recordings, CS: the number of control recordings with a spatial cell, LR: the number of lesion recordings, and LS: the number of control recordings with a spatial cell. As an example, using the data from the contingency table for the recordings with spatial cells (Table 1):

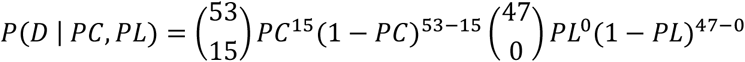

### Burst properties

Bursting units were identified using criteria based on Anderson and O’Mara^48^. A burst was defined as a series of spikes in which each inter-spike interval (ISI) was minimum of 6ms, and contained a minimum of two spikes, with a minimum inter-burst interval (IBI) of 50ms. Further bursting analyses examined the total number of bursts during a recording session; the number of spikes in the bursting cluster; mean inter-spike interval during the burst cluster; number of spikes per burst; burst duration; duty cycle (the portion of an inter-burst interval during which a burst fires); the inter-burst interval; and propensity to burst, calculated by dividing the number of bursting spikes by the total number of spikes in a recording.

### Spatial analyses

Additional analyses examined spatial modulation of recorded units. Multiple indices were used to analyse the spatial properties of unit activity (namely spatial coherence, spatial information content, and spatial sparsity). A firing field was defined as a set of at least nine contiguous pixels with firing rate above zero. A place field was identified if nine neighbouring pixels (sharing a side) were above 20% of the peak firing rate. Place field size was represented by number of pixels. Spatial specificity (spatial information content) was expressed in bits per spike^82^. Mean frequency is the total number of spikes divided by the total recording time and is expressed in Hz. Exploration was assessed by comparing the occupancy of bins and the number of visits per bin during recording sessions. Additionally, to be regarded as place cells, the following criteria had to be met: all included as place cells had to have a spatial information content^82^ index of > 0.5; a spatial coherence of > 0.25; and a mean firing rate > 0.25. The spatial path of the subject and the spike train were used to produce a locational firing rate map.

To analyse head direction (HD) cells, the animal’s head direction was calculated using the relative position of two tracked LEDs on a bar attached to the microdrive, in the horizontal plane. The directional tuning function was determined by plotting the firing rate as a function of the HD divided into 5° bins, and the firing rate calculated by the total number of spikes divided by time spent in each bin.

To determine the existence of a hexagonal grid firing structure, grid index, size and orientation were calculated. The grid cell analysis included calculating the spatial autocorrelation of the firing rate map and assessing the shape formed by the peaks in autocorrelation. For border cell analyses, cells with a firing profile that was parallel to the border of the arena were selected by plotting the positional firing pattern.

### Down-sampling analysis

We performed a spatial down-sampling procedure as a control method to rule out the possibility that the disruption in spatial firing merely reflects a lack of sampling of the environment, given that the animals move less distance and cover less of the environment while under the effects of muscimol (Fig. 6B).

Our method is based on the spatial down-sampling performed by Boccara et al.^83^: for each cell, we produce a list *L* = (*x_t, y_t, spike_t*) where *x_t, y_t* is the position of the animal at time *t* and spike_t is the number of spikes the cell emitted in that time bin. The procedure to spatially down-sample data from recording A to match the exploration in recording B is as follows. Firstly, the list L from B is binned into 3cm squares based on x_t, y_t and the list L from A is binned using the same number of bins as was used for B. Secondly, several random samples are drawn (with repetition) from the list L in A such that the number of samples drawn from an individual bin is the minimum of the number of data points falling in that bin between recordings A and B. Using this spatially down-sampled data, firing rate maps and spatial statistics are performed as in the rest of this paper. This procedure is performed 200 times for each cell.

Using the method above, recordings were spatially down-sampled to match their own spatial occupancy as an additional control since the random sampling procedure should cause small changes. After testing spatial coherence, spatial information content, and spatial sparsity, coherence was the only measure tested which was resistant to down-sampling against self (average decrease of 0.015) and down-sampling to other random data (average decrease of 0.05).

For each cell considered in the muscimol experiments, the baseline recording was down-sampled to match the muscimol recordings, and the muscimol recordings were down-sampled to match the baseline recording. These resulting firing maps had closely matching occupancy of the environment, and coherence was computed on these maps.

### Image analysis

Individual images of sections were aligned and tiled using Inkscape (http://inkscape.org) and FIJI (**F**iji **I**s **J**ust **I**mage J, https://imagej.net/Fij^79^). Cresyl violet stained sections helped to confirm electrode placement. In order to assess lesion success, the ATN was segmented from photomicrographs of anti-NeuN reacted sections using FIJI and the resulting image thresholded to show nuclei separation, before using the inbuilt Analyse Particles plugin to obtain a cell count. The ATN cell counts were compared between the ATNx and control groups to determine lesion effectiveness. Calbindin-reacted sections helped to determine the status of nucleus reuniens.

### Statistical analysis

Statistical analyses were performed in R Statistics (https://www.r-project.org/). No statistical difference was noted between Normal Control and Sham Control groups so these were combined into CControl unless otherwise stated. Differences between groups were examined using Welch’s Two-Sample *t*-test, an adaptation of Student’s *t*-test that is more reliable when samples have unequal variances or sample sizes. Where three groups were compared, ANOVA with a Tukey post-hoc test was applied.

For temporary inactivation experiments, data were compared to baseline (immediately prior to muscimol infusion) values using Welch’s Two-Sample *t*-test, with a Bonferroni correction to account for multiple comparisons and reduce the likelihood of Type 1 errors.

Statistical tests are reported in the text and appropriate figure legends, (* p < 0.05, ** p < 0.01, *** p < 0.001). Boxplots show median, 25^th^ and 75^th^ percentiles. Boxplot tails represent the smallest and largest value within 1.5 times the interquartile range, and outliers are defined as values that are > 1.5 times and < 3 times the interquartile range. Data means are represented with a diamond-shaped marker. Error bars on scatter plots represent standard error of the mean (SEM).

## Data and Code Availability

Datasets generated during this study are available at OSF (https://osf.io/vdakx/) and code is available at GitHub (https://github.com/shanemomara/omaraneurolab). Further information will be available upon request by contacting Shane O’Mara.

## Supplemental Information

**Supplementary Table 1:** Summary of spatial properties of subicular units before and 40-60 minutes after muscimol infusion. Cells which were recorded at baseline, prior to infusion, are included, although not all cells were recorded throughout the experiment or recorded the next day.

|  | Baseline |  | 40-60 min post muscimol infusion |  | Next Day |  |
| --- | --- | --- | --- | --- | --- | --- |
|  | Mean | ± SD | Mean | ± SD | Mean | ± SD |
| **Place cells (n=3)** | (n=3) |  | (n=3) |  | (n=2) |  |
| Spiking frequency (Hz) | 1.09 | 0.26 | 0.59 | 0.30 | 1.35 | 0.46 |
| Spatial information content | 1.11 | 0.23 | 1.40 | 0.85 | 1.77 | 0.17 |
| Spatial sparsity | 0.31 | 0.05 | 0.28 | 0.18 | 0.17 | 0.02 |
| Spatial coherence | 0.44 | 0.13 | 0.34 | 0.05 | 0.48 | 0.11 |
| **Head direction cells (n=2)** | (n=2) |  | (n=2) |  | (n=1) |  |
| Spiking frequency (Hz) | 0.66 | 0.44 | 0.49 | 0.22 | 0.47 | n/a |
| HD information content | 0.69 | 0.47 | 0.31 | 0.01 | 1.67 | n/a |
| Cell 1 mean HD | 22.92° |  | 90.56° |  | n/a |  |
| Cell 2 mean HD | 76.65° |  | 132.67° |  | 72.84° |  |
| **Grid cells (n=5)** | (n=5) |  | (n=3) |  | (n=2) |  |
| Spiking frequency (Hz) | 2.32 | 1.77 | 1.27 | 1.03 | 1.44 | 1.50 |
| Grid spacing | 35.43 | 5.60 | 27.36 | 4.30 | 28.98 | 10.87 |
| Grid score | 1.00 | 0.32 | 1.36 | 0.09 | 0.91 | 0.47 |
| Grid orientation | 39.14 | 29.99 | 69.64 | 49.76 | 36.05 | 6.04 |
| **Border cells (n=2)** | (n=2) |  | (n=2) |  | (n=0) |  |
| Spiking frequency (Hz) | 2.50 | 1.00 | 1.90 | 1.70 | n/a | n/a |
| Border information content | 0.057 | 0.052 | 0.094 | 0.085 | n/a | n/a |

**Supplementary Table 2:**
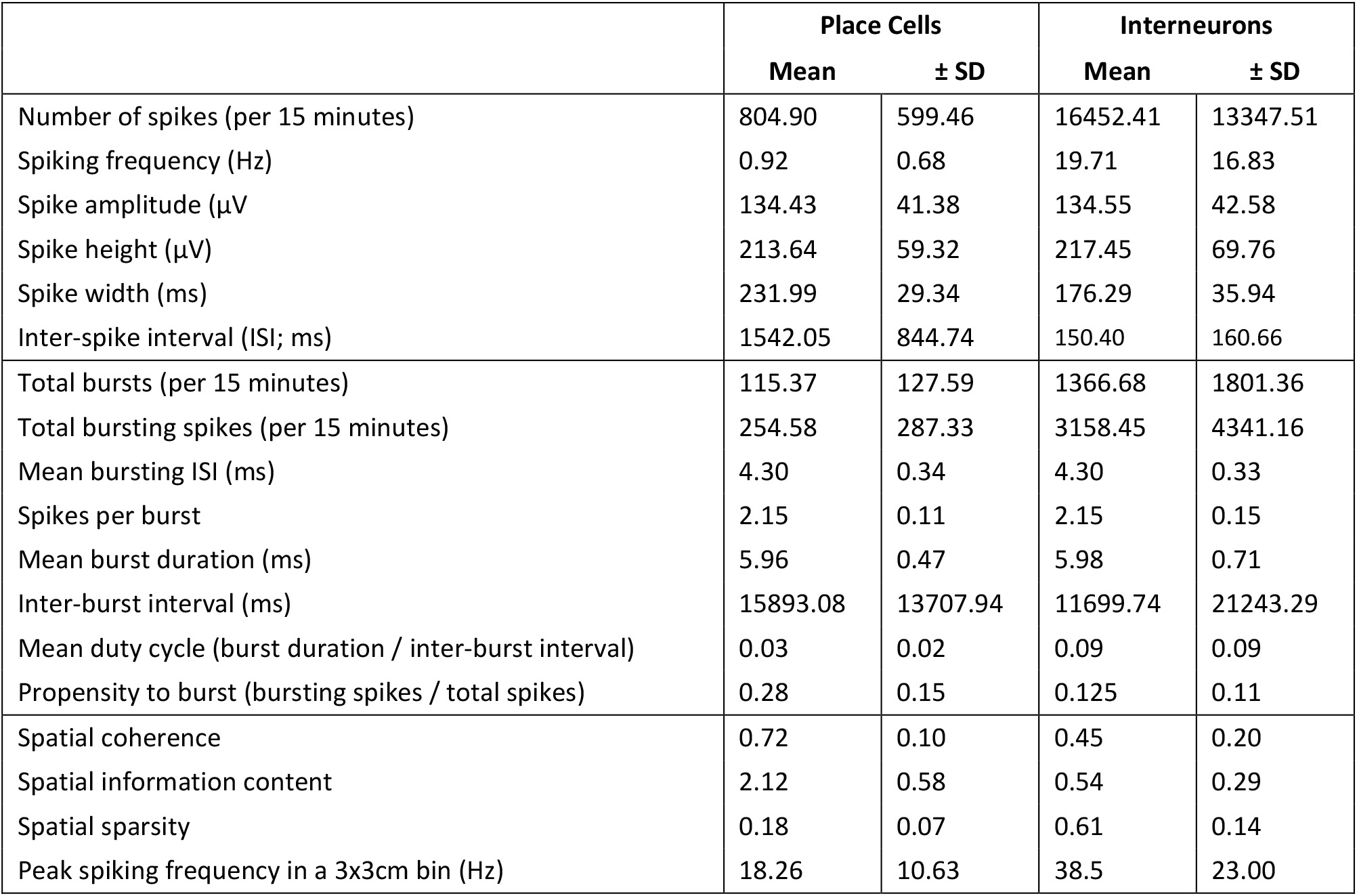
Summary of firing properties, burst properties, and spatial properties of dorsal CA1 units recorded in animals with ATN lesions. Cells were classified as pyramidal cells and interneurons based on their spike duration, waveform, firing frequency, and firing pattern^84^. Interneurons were only considered if they were present in the same recording as a pyramidal cell, as they were then CA1 pyramidal layer interneurons.

|  | Place Cells |  | Interneurons |  |
| --- | --- | --- | --- | --- |
|  | Mean | ± SD | Mean | ± SD |
| Number of spikes (per 15 minutes) | 804.90 | 599.46 | 16452.41 | 13347.51 |
| Spiking frequency (Hz) | 0.92 | 0.68 | 19.71 | 16.83 |
| Spike amplitude (μV) | 134.43 | 41.38 | 134.55 | 42.58 |
| Spike height (μV) | 213.64 | 59.32 | 217.45 | 69.76 |
| Spike width (ms) | 231.99 | 29.34 | 176.29 | 35.94 |
| Inter-spike interval (ISI; ms) | 1542.05 | 844.74 | 150.40 | 160.66 |
| Total bursts (per 15 minutes) | 115.37 | 127.59 | 1366.68 | 1801.36 |
| Total bursting spikes (per 15 minutes) | 254.58 | 287.33 | 3158.45 | 4341.16 |
| Mean bursting ISI (ms) | 4.30 | 0.34 | 4.30 | 0.33 |
| Spikes per burst | 2.15 | 0.11 | 2.15 | 0.15 |
| Mean burst duration (ms) | 5.96 | 0.47 | 5.98 | 0.71 |
| Inter-burst interval (ms) | 15893.08 | 13707.94 | 11699.74 | 21243.29 |
| Mean duty cycle (burst duration / inter-burst interval) | 0.03 | 0.02 | 0.09 | 0.09 |
| Propensity to burst (bursting spikes / total spikes) | 0.28 | 0.15 | 0.125 | 0.11 |
| Spatial coherence | 0.72 | 0.10 | 0.45 | 0.20 |
| Spatial information content | 2.12 | 0.58 | 0.54 | 0.29 |
| Spatial sparsity | 0.18 | 0.07 | 0.61 | 0.14 |
| Peak spiking frequency in a 3x3cm bin (Hz) | 18.26 | 10.63 | 38.5 | 23.00 |

**Supplementary Fig. 1:**
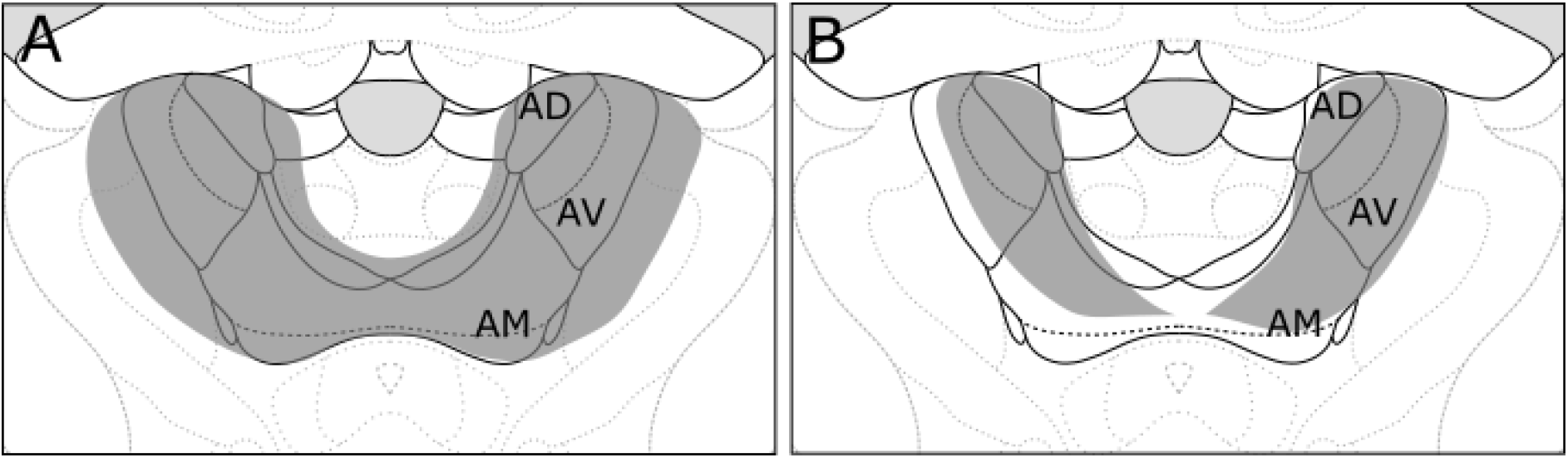
Graphical image showing maximum (A) and minimum (B) extent of NMDA lesion in ATNx-CA1 (grey).

**Supplementary Fig. 2:**
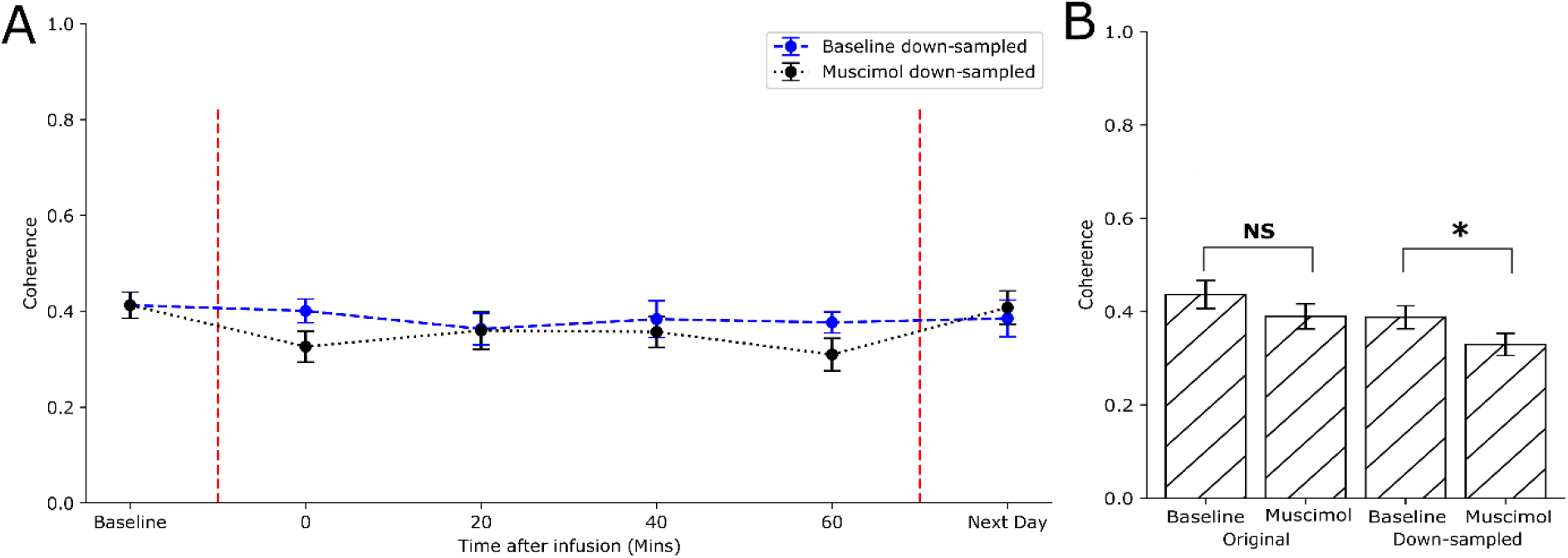
Results from spatially-down-sampling the subicular spatial units considered in the temporary muscimol ATN lesion experiments. (A) Comparison of average coherence over time in spatially down-sampled baseline recordings (blue line) and spatially down-sampled recordings performed after muscimol infusion (black line). Error bars represent standard error of the mean. (B) Bar chart representing the difference in coherence with and without down-sampling. The bars on the left show the average coherence values from baseline recordings of subicular spatial units and muscimol recordings without any down-sampling. The bars on the right show the average coherence values after the down-sampling procedure. NS indicates p >= 0.05, * indicates p < 0.05 (Welch’s Two Sample *t*-test).

